# Multidimensional *in vitro* assay for antimalarial combination testing and pharmacodynamic modeling – the MULT-i^2^ assay

**DOI:** 10.64898/2026.05.06.723157

**Authors:** Angela Hellingman, Christin Gumpp, Jörg J. Möhrle, Belen Tornesi, Didier Leroy, Sergio Wittlin, Pascal Mäser, Nicolas M. B. Brancucci, Sebastian G. Wicha, Matthias Rottmann

**Author notes:** Corresponding authors: Matthias Rottmann, Sebastian G. Wicha and Nicolas M.B. Brancucci.

## Abstract

Malaria remains a major global health challenge, with emerging partial resistance to first-line therapies in Africa threatening current control efforts. Drug combinations are essential to improve treatment efficacy and restrain resistance development. However, *in vitro* assays that quantify parasite viability after drug exposure and characterize pharmacodynamic drug interactions are labor- and resource-intensive, with standard approaches such as the parasite reduction ratio assay limiting systematic, high-resolution evaluation of drug combinations.

We present the *MU*ltidimensional *L*uminescence *T*est for *i*ntegration of *i*nteractions (MULT-i^2^), an *in vitro* assay that enables scalable, high-resolution assessment of parasite viability across multidimensional drug concentration spaces. For dual drug combinations, the MULT-i^2^ assay characterizes interaction surfaces while requiring ∼50-fold fewer resources and more than two-fold less time than conventional methods, enabling exploration of broader combination scenarios. The assay combines a highly sensitive chemiluminescence readout with inducible reporter expression in *Plasmodium falciparum,* supporting potential extension to multidimensional combination testing.

Using the general pharmacodynamic interaction (GPDI) model, the MULT-i^2^ assay quantified interaction potency and directionality, confirming and refining the known synergy between atovaquone and proguanil, and revealing detailed interaction patterns for additional drug combinations. Overall, this approach provides an efficient framework for testing and characterizing pharmacodynamic drug interactions and supports the rational development of antimalarial combination therapies.

## Introduction

Malaria is an infectious disease that continues to pose a major global health challenge, with more than 250 million cases and around 610’000 deaths in 2024 (Daily & Parikh, 2025; Maji, 2018; Sinha et al., 2017; World Health Organization, 2025). Artemisinin-based combination therapies (ACTs), the current first-line treatment for malaria, are increasingly threatened by the emergence of partial resistance to artemisinin and its derivatives, as well as rising resistance to their partner drugs (Dhorda et al., 2021; Phyo et al., 2016; van der Pluijm et al., 2019). Therefore, novel drug candidates and optimized treatment regimens are required to maintain effective therapies against *Plasmodium* infections and reduce the risk of resistance emergence (Blasco et al., 2017; Conrad et al., 2023; Dhorda et al., 2021; Menard & Dondorp, 2017; Rosenthal et al., 2024; World Health Organization, 2025).

*In vitro* antimalarial drug screening assays are essential tools for discovering and characterizing novel drug candidates against malaria (Daily & Parikh, 2025; Maji, 2018; Sinha et al., 2017; World Health Organization, 2025). Conventional growth-inhibition assays have been used for characterization of potential drug candidates, but they cannot distinguish between quiescent and dead parasites. As a result, isobolograms – used to evaluate drug-drug interactions and based on growth-inhibition assays – also fail to provide information on parasite-killing interactions of tested drugs (de Carvalho et al., 2023; Maiga et al., 2025; Sanz et al., 2012; Walz et al., 2023; Wicha et al., 2022). Parasite viability, regarded as the superior measure for antimalarial drug activity (Radohery et al., 2022; Rebelo et al., 2021), is a crucial parameter for pharmacometric modeling and for translating drug interactions into *in vivo* settings. To quantify *in vitro* parasite viability after drug exposure, the parasite reduction ratio (PRR) assay was developed (de Carvalho et al., 2023; Sanz et al., 2012; Walz et al., 2023). This assay allows to quantify key pharmacodynamic parameters such as the parasite clearance time (PCT), the PRR, the half-maximal effective concentration (EC50), and the maximum effect (Emax) from resulting time-kill profiles. An extension of the PRR assay, the combination PRR (cPRR) assay, enables the testing of drugs in combinations (Wicha et al., 2022). Results can then be used to model pharmacodynamic drug interactions and quantify directionality and intensity of these interactions (Wicha et al., 2022).

One major limitation of the PRR and cPRR assay is its material- and labor-intensive setup. The assay requires serial dilution of parasites and an extensive regrowth period for sensitive detection of viable parasites via conventional readouts such as [^3^H]-hypoxanthine incorporation (Sanz et al., 2012; Walz et al., 2023) or histidine-rich protein 2 enzyme-linked immunosorbent assays (HRP-2-ELISA) (de Carvalho et al., 2023). An optimized PRR assay using chemiluminescence-based readout could reduce workload and recovery time, nevertheless parasite serial dilution remains necessary (Hellingman et al., 2026). Therefore, systematic *in vitro* testing of dual or triple drug combinations is therefore extremely limited, increasing the risk of missing im portant pharmacodynamic interactions due to the restricted interaction surface tested. Novel assays based on direct viability assessment using MitoTracker dyes in combination with nuclear staining allow higher throughput viability testing without serial parasite dilution, but rely on flow cytometry-based readout, which also limits applicability for large-scale dual or triple combination testing (Maiga et al., 2024; Maiga et al., 2025). Whether the staining with MitoTracker can be considered a reliable direct marker of parasite viability, without confirmation of post-treatment recovery, still requires careful evaluation. Because MitoTracker dyes can accumulate in other organelles that possess a membrane potential but are not markers of cell viability, it could potentially generate artefactual labeling (Neikirk et al., 2023). When the analysis is based solely on flow cytometry, such signals cannot be readily distinguished from true mitochondrial staining, creating the possibility of misclassifying non-viable parasites as viable. To overcome current limitations in drug interaction testing, alternative approaches are required.

Here we developed an assay for systematic antimalarial drug interaction testing and modeling. Therefore, we combined a transgenic *P. falciparum* line with highly sensitive next generation chemiluminescent probes (Green et al., 2017; Hananya et al., 2016; Hananya & Shabat, 2017, 2019; Hellingman et al., 2024). To this end, we engineered *P. falciparum* parasites that allow for a conditional reporter enzyme production, serving as a highly sensitive marker to detect viable parasites following drug exposure. In a second step, we used this parasite line to establish the novel *MU*ltidimensional *L*uminescence *T*est for *i*ntegration of *i*nteractions (MULT-i^2^) assay. The resulting assay data were used to quantify pharmacodynamic parameters (e.g., EC50 and Emax) and interaction parameters (e.g., interaction intensity and potency) for antimalarial compounds and their combinations using the general pharmacodynamic interaction (GPDI) model (Wicha et al., 2017).

## Results

### Engineering of the inducible lacZ expression line NF54^i-lacZ^

The chemiluminescent β-gal^SENSOR^ probe (AquaSpark β-D-galactoside, cat. #A-8169_P00, Biosynth AG, Staad, Switzerland) can be activated by β-galactosidase encoded by the *lacZ* gene (Green et al., 2017; Hananya et al., 2016). We previously engineered transgenic *P. falciparum* parasites expressing a codon optimized *Escherichia coli lacZ* (Hellingman et al., 2024). To minimize background luminescence and to exclusively quantify viable parasites, we engineered a *P. falciparum* line carrying an inducible *lacZ* expression reporter. In these *P. falciparum* NF54*^i-lacZ^*, exposure to rapamycin (Rapa) induces heterodimerization and activation of the Cre recombinase, which then selectively recognizes the two loxP sites we introduced into the *P230p* locus (PF3D7_0208900) of NF54^DiCre^ parasites (Figure 1A and 1B) (Tibúrcio et al., 2019). Specifically, we inserted an expression cassette in which *green fluorescent protein (gfp)* expression is controlled by the stress-response promoter of *heat shock protein 70 (hsp70)* (Bianco et al., 1986; Haas, 1994; Przyborski et al., 2015) and a *calmodulin (cam)* terminator (Ashdown et al., 2020; Collins et al., 2013; Ghorbal et al., 2014; Jones et al., 2016; Tibúrcio et al., 2019). The loxP-sites are situated within an intron (loxPint) (Das et al., 2015; Jones et al., 2016) at the 5’-end of the *gfp* coding sequence as well as directly downstream of the *cam* terminator. The genome editing strategy used to generate NF54*^i-lacZ^* is displayed in Figure 1 – figure supplement 1. Upon rapamycin dependent DiCre activation, the *gfp* coding sequence is disrupted and *lacZ* is placed under control of the *hsp70* promoter instead. This renders *lacZ* expression – and consequently chemiluminescence-based detection via enzymatic activation of the β-gal^SENSOR^ probe – inducible by rapamycin.

**Figure 1:**
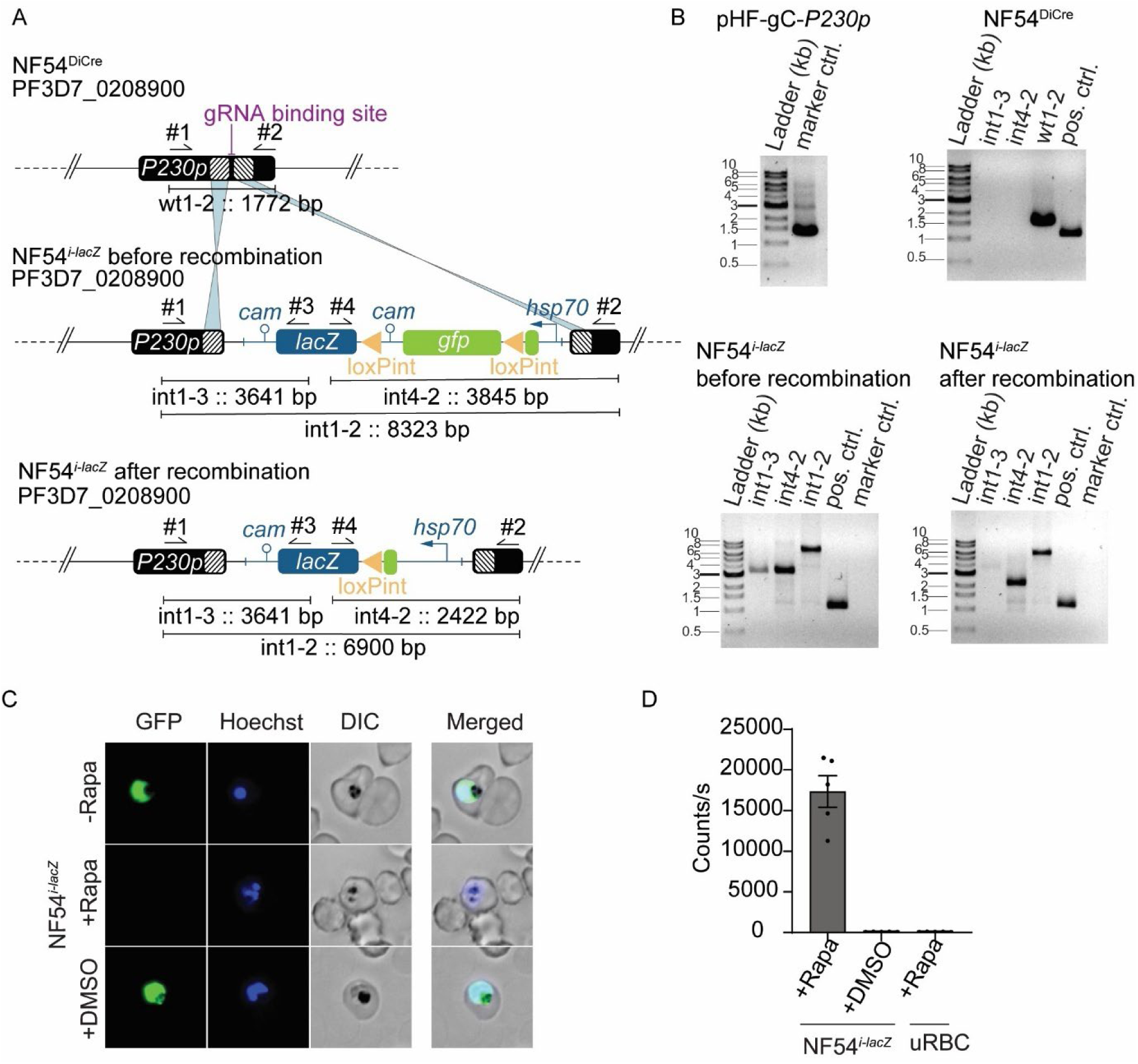
Inducible *lacZ* expression in the *P. falciparum* NF54*^i-lacZ^.* (A) Schematic representation of the *P230p* locus in parental NF54^DiCre^ and NF54*^i-lacZ^* parasites before and after rapamycin-induced recombination. Arrows indicate primers used for diagnostic integrations PCRs. Striped boxes represent homology regions used for CRISPR/Cas9-based gene editing. The gRNA binding site is indicated. loxPint, loxP-intron. (B) Diagnostic PCR confirmed the integration of the loxPInt-*gfp-lacZ* expression cassette in *P. falciparum* NF54*^i-lacZ^,* and DiCre-induced recombination. Diagnostic *hdhfr-yfcu* (1595 bases) PCR demonstrates loss of selectable markers in the NF54*^i-lacZ^* (marker control). Primers used are shown in Figure 1A and Figure 1 – figure supplement 1 (#1-#6). wt: wild type; int: integration; pos. ctrl: positive control (1252 bp). (C) GFP and Hoechst fluorescence signals, and (D) chemiluminescence emitted from NF54*^i-lacZ^* parasites before (-Rapa) or after (+Rapa, initial parasitemia of 0.3% measured after 48 h) rapamycin-induced recombination, and controls (vehicle control (+DMSO), uRBC control treated with Rapa); error bars represent the standard error of the mean (SEM) of 5 biological replicates conducted in at least technical duplicates. Rapa: rapamycin; GFP: green fluorescent protein; DIC: differential interference contrast; uRBC: uninfected red blood cells. The β-gal^SENSOR^ probe was used at 10 µM and rapamycin at 100 nM. **Figure 1B – source data 1:** Original agarose gel image of diagnostic PCRs confirming the integration of the loxPInt-*gfp*-*lacZ* expression cassette in *P. falciparum* NF54*^i-lacZ^*, successful recombination after DiCre-induced recombination, and loss of selectable markers in the NF54*^i-lacZ^* (marker control). **Figure 1D – source data 2:** Chemiluminescence signal after rapamycin-induced recombination measured 48 hours after rapamycin addition. **Figure 1 – figure supplement 1:** Overview of the two-plasmid CRISPR/Cas9-based gene editing strategy. Schematic of the *P230p* locus in NF54^DiCre^ parasites and the plasmids used for CRISPR/Cas9-mediated gene editing (*P230p*-loxPInt-gfp-*lacZ* and pHF-gC-*P230p*) to generate the NF54*^i-lacZ^* parasites. **Figure 1 – figure supplement 2:** Susceptibility to the antifolate drug WR99210 and parasitized erythrocyte infection rate of the novel NF54^i-lacZ^ compared to NF54^WT^. (A) IC50 values of WR99210; error bars represent the standard error of the mean (SEM) of three biological replicates conducted in technical duplicates and (B) parasitized erythrocyte infection rates (within a period of 48 hours) are comparable between NF54^i-lacZ^ parasites, either rapamycin-treated or untreated, and the NF54^WT^ strain; error bars represent the SEM of three biological replicates. Rapa: rapamycin. **Figure 1 – figure supplement 2 – source data 1:** Measured and normalized parasite growth (% of NF54^wt^ or NF54*^i-lacZ^*) versus WR99210 concentration (nM) used for IC50 determination and inhibitory dose–response curve fitting. **Figure 1 – figure supplement 2 – source data 2:** Microscopically counted parasitemia and proportion of ring stages (%) used to calculate the parasitized erythrocyte infection rate within 48 hours (one asexual intraerythrocytic developmental cycle).

As anticipated, *P. falciparum* NF54*^i-lacZ^* parasites exhibit robust *gfp* expression in the absence of rapamycin or following treatment with the dimethyl sulfoxide (DMSO) vehicle control. By contrast, *gfp* expression is completely abolished after 48 hours of rapamycin treatment (Figure 1C). Conversely, rapamycin-induced DiCre-mediated recombination activates *lacZ* transcription, resulting in the production of β-galactosidase, which subsequently cleaves and activates the β-gal^SENSOR^ probe, resulting in the emission of a luminescence (Figure 1D). Luminescence cannot be detected in parasites treated with the vehicle of rapamycin (DMSO) and is identical to that observed in rapamycin-treated uninfected red blood cells (uRBCs). Cre-mediated recombination in NF54*^i-lacZ^* is highly efficient and averaged 97% (SD: ∼0.4%, of 3 biological replicates). We further confirmed that the drug selectable marker used for transgene insertion (*hdhfr* on the pHF-gC-*P230p* plasmid) is absent in NF54*^i-lacZ^* using half-maximal inhibitory concentration (IC50) analysis of the antifolate drug WR99210 (Figure 1 – figure supplement 2A). The parasitized erythrocyte infection rate of NF54*^i-lacZ^*, treated with or without rapamycin, was comparable to that of the wild type NF54 strain (Figure 1 – figure supplement 2B). Unless stated otherwise, subsequent experiments were conducted using the *P. falciparum* NF54*^i-lacZ^* line.

### Limit of quantification of recombined P. falciparum NF54*i-lacZ*

Following rapamycin-induced DiCre-mediated recombination, parasites require time to produce detectable levels of the β-galactosidase reporter enzyme. To determine the limit of quantification (LOQ) for the chemiluminescent signal generated by the β-gal^SENSOR^ probe, we serially diluted asynchronous *P. falciparum* NF54*^i-lacZ^* parasites. The diluted parasites were incubated for varying periods in presence of rapamycin prior to quantifying chemiluminescence (Figure 2).

**Figure 2:**
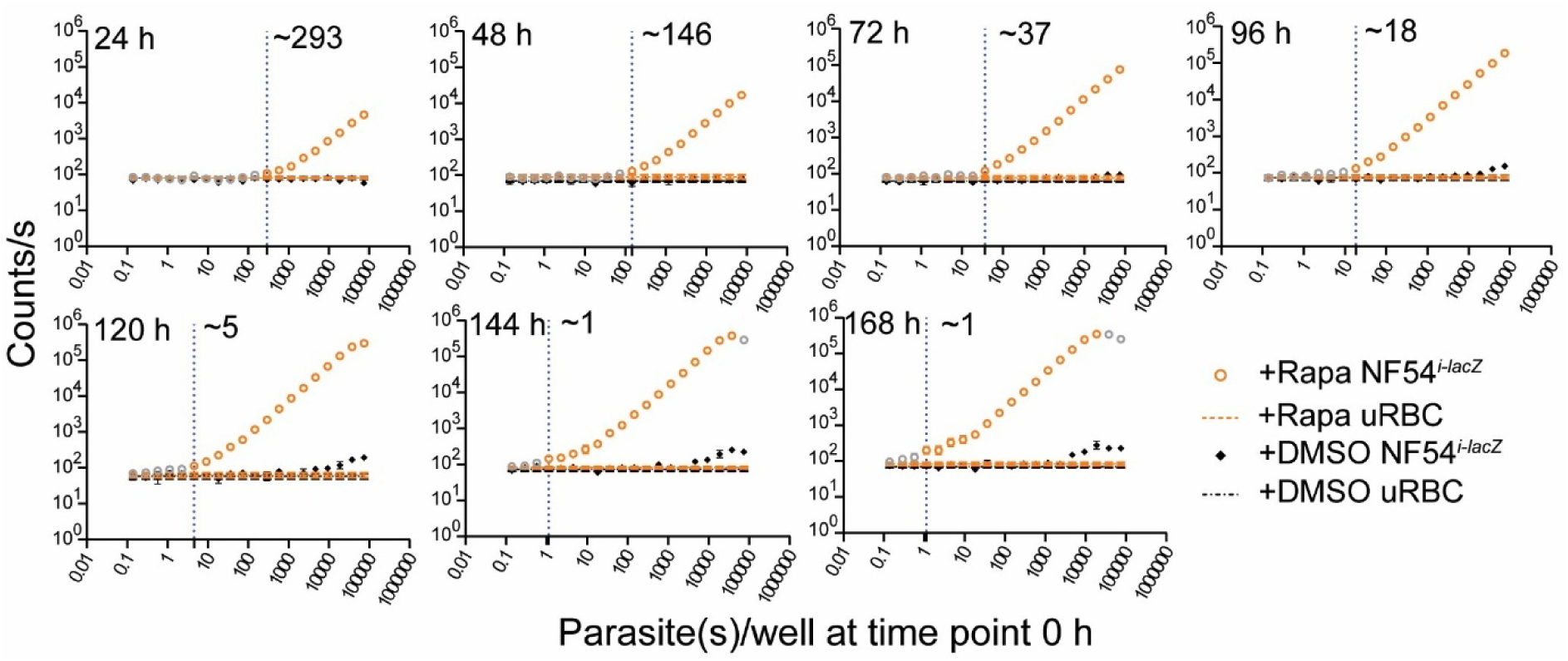
Limit of quantification of recombined *P. falciparum* NF54*^i-lacZ^* parasites. Serially diluted NF54*^i-lacZ^* parasites were incubated for varying durations (24-168 hours) with 100 nM rapamycin or the corresponding concentration of DMSO before chemiluminescence measurement using 10 μM β-gal^SENSOR^ probe. The signal linearly correlated with the initial parasite inoculum, improving the limit of quantification over time from ∼300 initial parasites per well to as low as ∼1 initial parasite(s) per well (indicated by the vertical dotted line). Controls (uRBC treated with rapamycin or DMSO) are indicated. The LOQ was tested in ≥ three biological replicates with at least technical duplicates; error bars represent the standard error of the mean (SEM) of biological replicates and may not be visible where smaller than the plotted symbols. The LOQ was set to the final initial parasite number per well for which the generated signal against the negative control (uRBC treated with Rapa) was considered significantly different in an unpaired t-test (p < 0.05) and a linear relationship was given. uRBC: uninfected red blood cells; Rapa: rapamycin. **Figure 2 – source data 1:** Measured chemiluminescence signal for the initial parasite inoculum or the uninfected red blood cell control after the corresponding incubation time under 100 nM rapamycin or DMSO.

The chemiluminescent signal exhibited a strong linear correlation with the initial parasite inoculum across a broad dynamic range. The LOQ improved progressively with longer incubation periods. Specifically, the LOQ corresponded to approximately 300 initial parasites per well after 24 hours of growth, decreased to ∼5 initial parasites per well after 120 hours, and reached ∼1 initial parasite per well after 144 hours or longer. At high initial parasite counts, the signal intensity plateaued or declined after ≥ 120 hours of incubation. In parallel, DMSO-treated control parasites displayed a gradual increase in background signal for high initial parasite inoculums after ≥ 96 hours. Balancing the gains in sensitivity of prolonged rapamycin exposure with practical considerations related to assay duration and laboratory workflow, we selected 120 hours of rapamycin incubation as the standardized condition for all subsequent experiments.

### The MULT-i^2^ assay reduces material, time and workload for viability testing of parasites after drug treatment

To enable quantification of viable parasites after drug exposure without the need for serial dilution, we established the MULT-i^2^ assay protocol, which is based on the previously reported *lacZ*/β-gal^SENSOR^ system (Hellingman et al., 2024) and utilizes the inducible *P. falciparum* NF54*^i-lacZ^*line. The inducible system prevents reporter accumulation during drug pressure, therefore reduces background signal by the stable reporter enzyme, and enables the quantification of living parasite numbers after drug exposure.

The MULT-i^2^ assay allows versatile time-kill assessments of single- and dual-drug combinations with the possibility to extend to triple-drug combinations. Figure 3 schematically illustrates the MULT-i^2^ assay protocol for dual-drug combination testing, which can be extended to triple combinations by adding a third dimension to the assay design. Dual drug combinations are prepared in a checkerboard format that includes all concentrations tested in monotherapy as well as all possible pairwise combinations (Bellio et al., 2021), and subsequently distributed into 96-well assay plates (Figure 3). Asynchronous *P. falciparum* NF54*^i-lacZ^* parasites are then added, and the plates are incubated for the desired duration under drug exposure. After incubation, the drug(s) are removed by several washing steps, and rapamycin is added to induce *lacZ* expression. Without the need to serially dilute the parasites, plates are then incubated for an additional 120 hours to allow surviving parasites to grow and accumulate β-galactosidase through recombination, gene expression, and protein synthesis. The plates are frozen prior to the readout, allowing flexible scheduling and convenient sample handling prior to measurement. At the start of each assay, a reference calibration “0-hour” plate is prepared in parallel using the same parasite culture as for the drug treated plates. Parasites in this calibration plate are serially diluted prior to 120 hours rapamycin treatment to generate a standard curve. Luminescence data can then be acquired together with the test plates and used for pharmacodynamic modeling.

**Figure 3:**
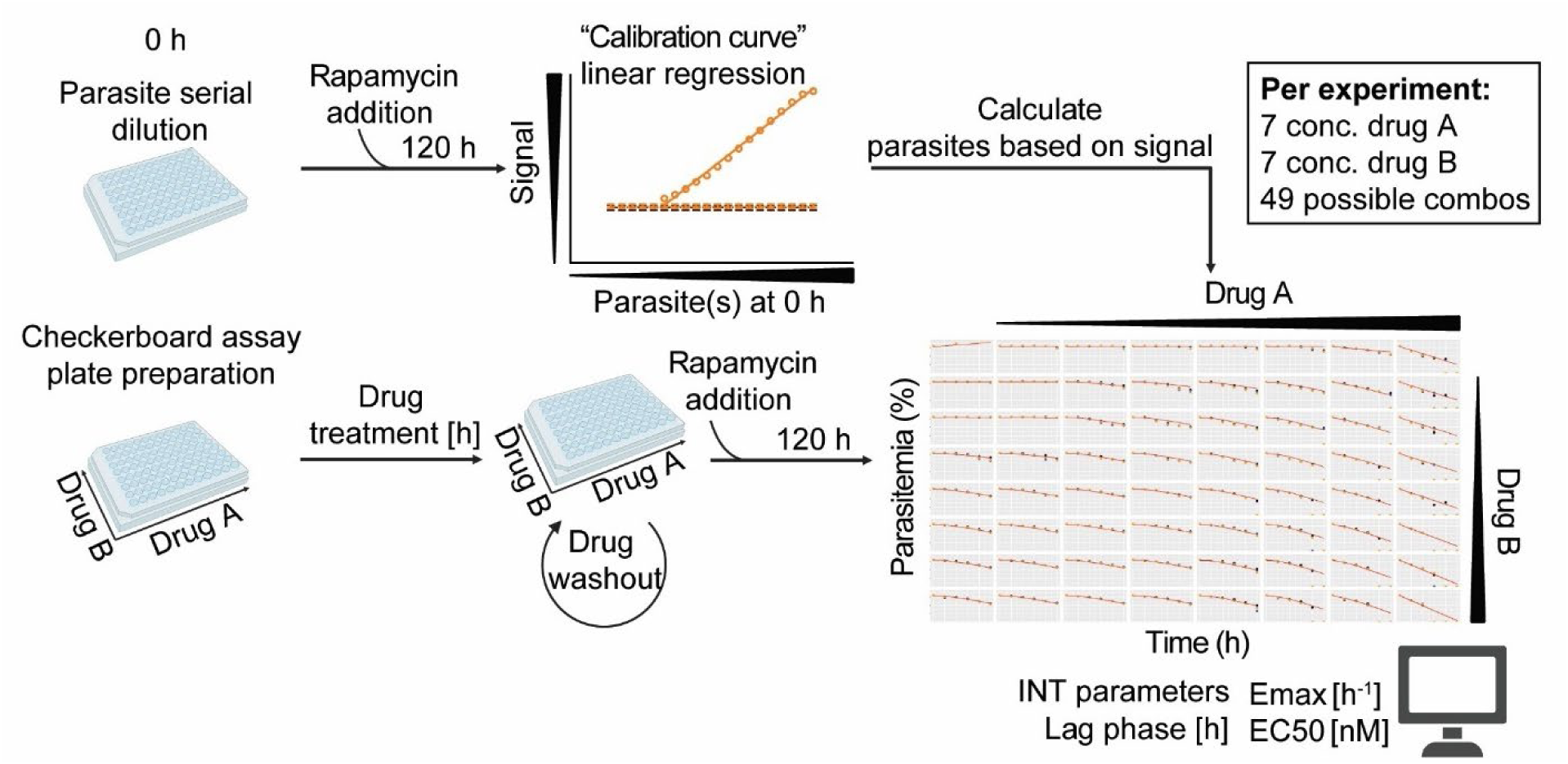
Schematic overview of the MULT-i^2^ assay workflow for dual drug combination testing. A 0 h calibration plate containing a parasite serial dilution, starting at 10⁵ parasites, is prepared as a reference. In parallel, checkerboard plates with the desired drug concentrations and combinations are assembled, and 10⁵ parasites are added per condition. The 0 h calibration plate is immediately treated with rapamycin and incubated for 120 h to induce *lacZ* expression and β-galactosidase accumulation. Drug-treated plates are incubated under drug pressure for the desired duration, after which drugs are removed by extensive washing. Rapamycin is then added for 120 h to enable surviving parasites to express the reporter enzyme β-galactosidase. Plates are subsequently frozen for later chemiluminescent signal measurement. The 0 h calibration plate establishes the relationship between chemiluminescence signal intensity and initial parasite number, allowing quantification of viable parasites after drug exposure. In this configuration, the assay supports testing of two compounds (A and B) across seven concentrations each, yielding 49 dual combinations. The resulting data are used to model pharmacodynamic parameters using the GPDI framework, including interaction parameters (INT), lag phases, maximal effect (Emax), and half-maximal effective concentration (EC50). Figure 3 was partly created with BioRender.com.

Using this novel MULT-i^2^ assay workflow, a single experiment testing a dual-drug combination (drugs A and B, each at seven single concentrations and all 49 possible dual combinations) across five time points can be completed within ∼11 days. This requires only six to seven 96-well plates and one 6-well plate. In contrast, performing the same experiment with the conventional combination PRR (cPRR) assay would require 21-28 days and around 380 assay plates (∼320×96- and ∼63×6-well plates), in addition to substantially higher manual workload. This represents a reduction in required resources of more than 50-fold and a decrease in overall assay duration of >2-fold. This improvement applies to single-drug testing as well as triple-combination testing.

### Time-killing curves of reference compounds generated with the MULT-i^2^ assay are comparable to curves generated with the PRR v2

The MULT-i^2^ assay protocol was validated using the *P. falciparum* NF54*^i-lacZ^* strain by testing four reference compounds – artemisinin, chloroquine, pyrimethamine, and atovaquone – which represent drugs with distinct modes and speeds of action. We then compared these results with those published by Walz et al., who developed and used the PRR version 2 (PRR v2) protocol in combination with wild type NF54 parasites (Walz et al., 2023). Therefore, we performed the MULT-i^2^ assay using the same compound concentrations as in the PRR v2 and analyzed the resulting data with the R pipeline described by Walz et al. (Walz et al., 2023).

Overall, the time-killing profiles of the tested drugs were consistent between both assays, with artemisinin showing the fastest parasite clearance and atovaquone the slowest (Figure 4). For fast-acting compounds such as artemisinin and chloroquine, the pharmacodynamic parameters aligned very closely between the two assays. For pyrimethamine, a slower acting compound, the new MULT-i^2^ assay predicted smaller log_10_(PRR) of 2.7 compared to 3.8 with the PRR v2 assay (Table 1). The largest discrepancy was detected for atovaquone with a shorter predicted lag time of 12 h compared to the previously reported 48 h. Overall, both assays showed comparable time-killing profiles for the four reference antimalarials, and the MULT-i^2^ assay detected viable parasites in all conditions where the PRR v2 assay did.

**Figure 4:**
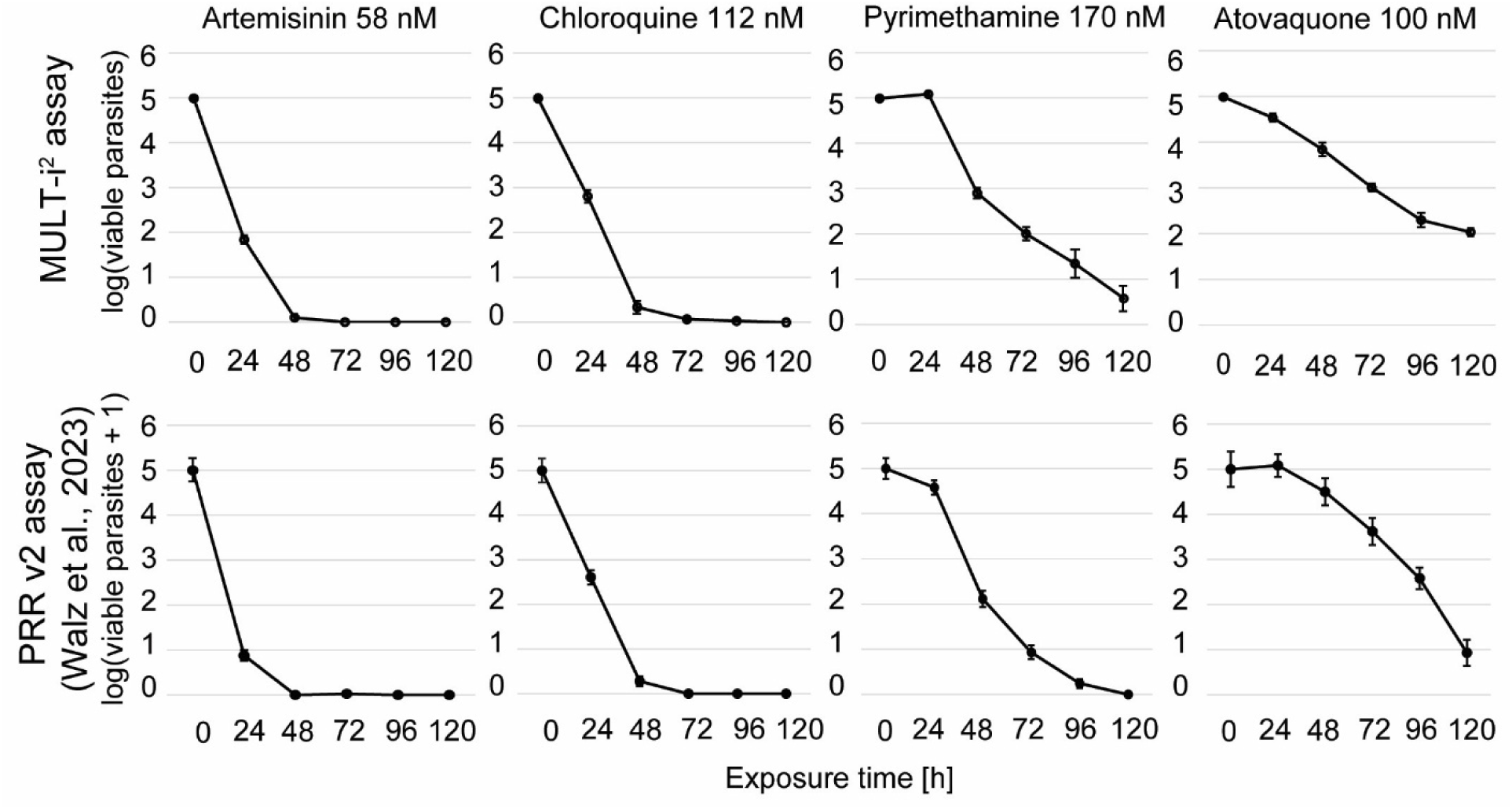
Comparison of time-killing profiles of four reference compounds between the MULT-i^2^ assay and the published PRR v2. Artemisinin, chloroquine, pyrimethamine and atovaquone were tested in the MULT-i^2^ assay using the same compound concentration as in the published PRR v2 (Walz et al., 2023). Data points represent the mean of ≥ three biological replicates (two for the 120-hour time point in the MULT-i^2^ assay) in four technical replicates, error bars represent the standard error of the mean (SEM) of the biological replicates. **Figure 4 – source data 1:** Dataset of the MULT-i^2^ assay containing the log(viable parasites) per tested time point of the four reference compounds used for plotting time killing profiles and calculating lag time, log10(PRR), PCT99.9% and Emax with the R pipeline published by Walz et al., 2023.

**Table 1:**
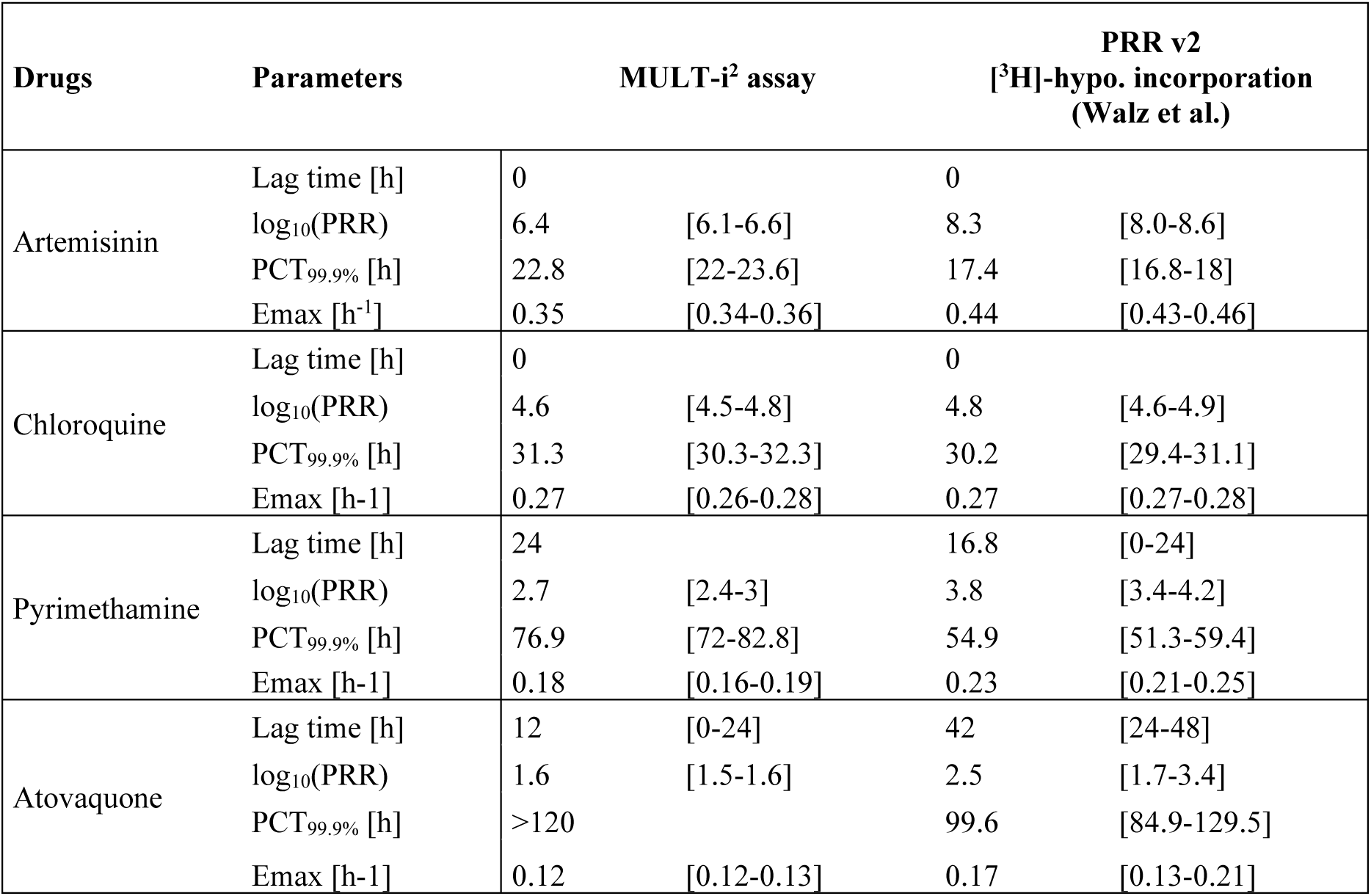
Comparison of pharmacodynamic parameters calculated with the published R pipeline (Walz et al., 2023), based on data from ≥ three biological replicates in one (MULT-i^2^ assay) in four technical replicates. Log_10_(PRR): log_10_(parasite reduction ratio); PCT_99.9%_: 99.9% parasite clearance time (h); Emax: maximal drug effect – parasite killing rate (h^−1^); hypo.: hypoxanthine. The 95% confidence interval is shown in square brackets. **Table 1 – source data 1:** same as indicated in Figure 4 – source data 1

| Drugs | Parameters | MULT-$i^2$ assay | | PRR v2<br>[<sup>3</sup>H]-hypo. incorporation<br>(Walz et al.) | |
| --- | --- | --- | --- | --- | --- |
| Artemisinin | Lag time [h] | 0 |  | 0 |  |
|  | log<sub>10</sub>(PRR) | 6.4 | [6.1-6.6] | 8.3 | [8.0-8.6] |
|  | PCT<sub>99.9%</sub> [h] | 22.8 | [22-23.6] | 17.4 | [16.8-18] |
|  | Emax [h<sup>-1</sup>] | 0.35 | [0.34-0.36] | 0.44 | [0.43-0.46] |
| Chloroquine | Lag time [h] | 0 |  | 0 |  |
|  | log<sub>10</sub>(PRR) | 4.6 | [4.5-4.8] | 4.8 | [4.6-4.9] |
|  | PCT<sub>99.9%</sub> [h] | 31.3 | [30.3-32.3] | 30.2 | [29.4-31.1] |
|  | Emax [h<sup>-1</sup>] | 0.27 | [0.26-0.28] | 0.27 | [0.27-0.28] |
| Pyrimethamine | Lag time [h] | 24 |  | 16.8 | [0-24] |
|  | log<sub>10</sub>(PRR) | 2.7 | [2.4-3] | 3.8 | [3.4-4.2] |
|  | PCT<sub>99.9%</sub> [h] | 76.9 | [72-82.8] | 54.9 | [51.3-59.4] |
|  | Emax [h<sup>-1</sup>] | 0.18 | [0.16-0.19] | 0.23 | [0.21-0.25] |
| Atovaquone | Lag time [h] | 12 | [0-24] | 42 | [24-48] |
|  | log<sub>10</sub>(PRR) | 1.6 | [1.5-1.6] | 2.5 | [1.7-3.4] |
|  | PCT<sub>99.9%</sub> [h] | >120 |  | 99.6 | [84.9-129.5] |
|  | Emax [h<sup>-1</sup>] | 0.12 | [0.12-0.13] | 0.17 | [0.13-0.21] |

### The MULT-i^2^ assay allows detailed pharmacodynamic parameter modeling

Given the promising performance of the single-drug testing with the four reference compounds, we investigated the well characterized synergism between atovaquone and proguanil (Looareesuwan et al., 1996; Srivastava Indresh & Vaidya Akhil, 1999) with the *in vitro* MULT-i^2^ assay. The results were compared to those obtained with the cPRR assay (Wicha et al., 2022). The MULT-i^2^ assay was performed testing five time points with seven single-drug concentrations for each compound and all 49 possible dual combinations. The cPRR was performed in a reduced design testing three time points, with four single-drug concentrations for each compound and only nine dual combinations. Time dependent time-killing profiles for mono drug and dual drug combinations were generated for both assays as shown in Figure 5.

**Figure 5:**
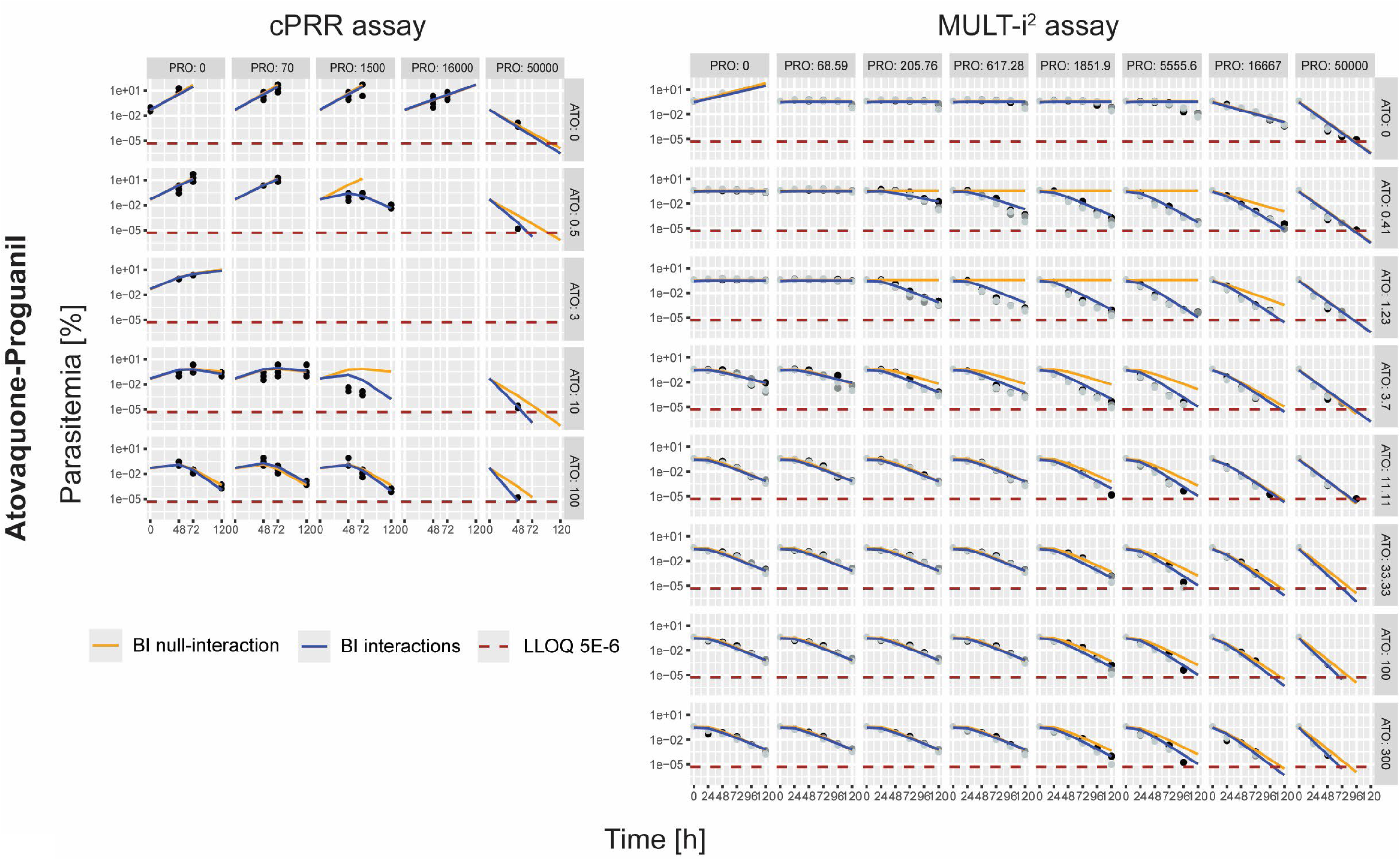
Model fits based on cPRR assay or MULT-i^2^ assay data for the tested combination atovaquone-proguanil. Model predictions are shown as orange (BI null-interaction) or blue (BI interaction) line, points represent original data (cPRR - one biological replicate with four technical replicates, three tested timepoints; MULT-i^2^ - three biological replicates with one technical replicate, five tested time points), the LLOQ is indicated by a dashed brown line. Dark grey boxes indicate the concentrations (nM) of the drugs used: ATO: atovaquone; PRO: proguanil; BI: Bliss Independence; LLOQ: lower limit of quantification. **Figure 5 – source data 1:** Original cPRR dataset used for modeling in the Nonlinear Mixed Effects Modelling (NONMEM) software containing measured values (also called dependent variables (DV), parasitemia %) at tested time points (TIME, hours), used drug concentrations for atovaquone (CA, nM) and proguanil (CB, nM). The original result datasets contain the final plotted model estimates (individual predicted values (IPRED), parasitemia %) for BI null-interaction (orange) or BI Interactions (blue). **Figure 5 – source data 2:** Original MULT-i^2^ dataset used for modeling in NONMEM containing measured values (DV, parasitemia %) at tested time points (TIME, hours), used drug concentrations for atovaquone (CA, nM) and proguanil (CB, nM). The original result datasets contain the final plotted model estimates (IPRED, parasitemia %) for BI null-interaction (orange) or BI Interactions (blue).

Using data from the MULT-i^2^ and cPRR assay, the null-interaction criterion Bliss Independence (Pearson et al., 2023), based on the single-drug effects of atovaquone and proguanil, underestimated the observed effects for certain combinations (Figure 5, orange line). A model incorporating a synergistic interaction between atovaquone and proguanil using the GPDI model (Wicha et al., 2017) provided a superior fit (Figure 5, blue line) than a model assuming null-interaction according to Bliss Independence (Δ Akaike Information Criterion (AIC) cPRR: 257.385; ΔAIC MULT-i^2^: 1962.608) for both assays. In the cPRR assay, synergistic interactions were observed at 0.5 and 10 nM atovaquone combined with 1500 nM and 50000 nM proguanil, which were also detected by the MULT-i^2^ assay. In addition, the MULT-i^2^ assay identified synergistic interactions at concentrations not accessible with the cPRR assay due to its minimal design (used due to resource limitations linked to the cPRR assay setup). The GPDI model using cPRR data identified three significant interactions: a symmetric synergistic EC50 shift of atovaquone by proguanil and vice versa, and a monodirectional antagonistic interaction on Emax. However, the Emax antagonism is not directly observable in the plotted data (Figure 5: null-interaction line (orange) lies above measured values) and is likely a model-driven artifact arising from limited information at high drug concentrations in the sparse data set. In contrast, the model using MULT-i^2^ data identified four significant interactions including asymmetric interaction on EC50 and symmetric synergism on Emax. Interaction classification on Emax differed between the cPRR and the MULT-i^2^ assay. Nevertheless, data from both approaches consistently identified the strongest interaction as a reduction in the EC50 of atovaquone (estimated EC50: 14.7 nM (cPRR) and 2.62 nM (MULT-i^2^)) by proguanil (reduction of 99% (cPRR) and 91.7% (MULT-i^2^)), with full parameter estimates provided in Supplementary Table 3.

To further validate the MULT-i^2^ assay in comparison with the cPRR, we tested the combination of the two fast acting drugs pyronaridine and piperaquine in a layout comparable to that used above. For this combination the GPDI model (Figure 6, the orange line representing Bliss Independence null-interaction, and the blue line resembling Bliss Independence with interactions) revealed synergism at concentrations around the EC50 (estimated EC50 of pyronaridine: 9.85 nM (cPRR) and 13.2 nM (MULT-i^2^); estimated EC50 of piperaquine: 25.5 nM (cPRR) and 25.8 nM (MULT-i^2^)), which converts to antagonism at high concentrations of both drugs and was detected in both assay types. The cPRR assay identified only two significant interactions: a monodirectional synergistic interaction on EC50 and a monodirectional antagonistic interaction on Emax, although the antagonism estimate was based on the limited datapoints available from the three combination scenarios using the highest tested pyronaridine concentration of 25 nM. In contrast, the MULT-i^2^ assay identified four significant interactions, comprising symmetric synergism on EC50 and symmetric antagonism on Emax. In this case, the antagonism observed at high drug concentrations was more evident, supported by additional data points across a broader concentration range, reaching up to 50 nM for pyronaridine and 200 nM for piperaquine. Also, for pyronaridine-piperaquine, the models considering interactions provided a superior fit compared to the models assuming null-interaction (ΔAIC cPRR: 6490.863; ΔAIC MULT-i^2^: 49’637.17). The two strongest interactions detected with both assays were the synergistic reduction of the EC50 of pyronaridine by piperaquine (reduction of 85% (cPRR) and 96.1% (MULT-i^2^), and the antagonistic effect of decreasing the Emax of pyronaridine by piperaquine (decrease by 99% (cPRR) and by 55.9% (MULT-i^2^). For both tested drug combinations, the relative standard errors of the estimated parameters and the residual errors, were smaller with the rich MULT-i^2^ assay dataset than with the cPRR (Supplementary Table 3).

**Figure 6:**
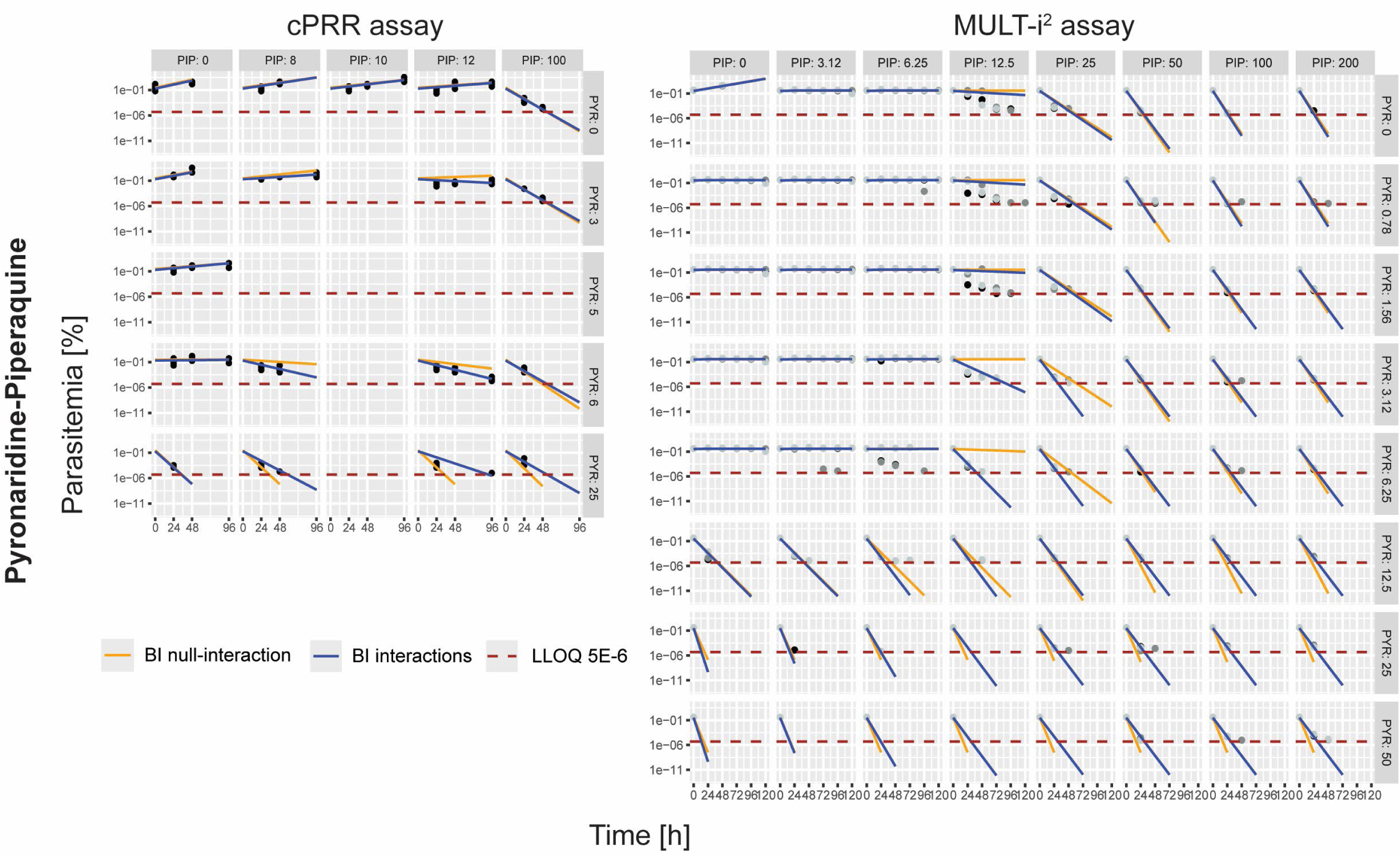
Model fits based on cPRR assay or MULT-i^2^ assay data for the tested combination, pyronaridine-piperaquine. Model predictions are shown as orange (BI null-interaction) or blue (BI interaction) line, points represent original data (cPRR - one biological replicate with four technical replicates, three tested time points; MULT-i^2^ - three biological replicates with one technical replicate, five tested time points), the LLOQ is indicated by a dashed brown line. Dark grey boxes indicate the concentrations (nM) of the drugs used: PYR: pyronaridine; PIP: piperaquine; BI, Bliss Independence; LLOQ, lower limit of quantification. **Figure 6 – source data 1:** Original cPRR dataset used for modeling in NONMEM containing measured values (DV, parasitemia %) at tested time points (TIME, hours), used drug concentrations for pyronaridine (CA, nM) and piperaquine (CB, nM). The original result datasets contain the final plotted model estimates (IPRED, parasitemia %) for BI null-interaction (orange) or BI Interactions (blue). **Figure 6 – source data 2:** Original MULT-i^2^ dataset used for modeling in NONMEM containing measured values (DV, parasitemia %) at tested time points (TIME, hours), used drug concentrations for pyronaridine (CA, nM) and piperaquine (CB, nM). The original result datasets contain the final plotted model estimates (IPRED, parasitemia %) for BI null-interaction (orange) or BI Interactions (blue).

## Discussion and Conclusion

Parasite viability is a superior measure for assessing the antimalarial activity of drug candidates (Radohery et al., 2022; Rebelo et al., 2021). The standard method to assess parasite viability after drug exposure, the parasite reduction ratio assay (PRR), is both material- and labor-intensive. As a result, it is poorly suited for studies that require testing large numbers of compounds, broad concentration ranges, or systematic dual- and triple-drug combinations (Sanz et al., 2012; Walz et al., 2023).

To address these limitations, we developed the scalable *in vitro* multidimensional luminescence test for integration of interactions (MULT-i^2^) assay for antimalarial combination testing. This assay combines next generation chemiluminescent probes (β-gal^SENSOR^ probe) (Green et al., 2017; Hananya et al., 2016; Hananya & Shabat, 2017, 2019) with an inducible *lacZ*-based reporter system. In the engineered *P. falciparum* NF54*^i-lacZ^*strain, rapamycin-induced recombination triggers *lacZ* expression encoding the reporter enzyme β-galactosidase, which activates the chemiluminescent β-gal^SENSOR^ probe. This method enables sensitive and efficient quantification of viable parasites without the labor-intensive serial dilution steps required in the PRR assay to extrapolate viable parasite numbers from the dilution factor, and positive parasite growth titers. The MULT-i^2^ assay leverages the linear correlation between β-gal^SENSOR^-derived chemiluminescence and initial parasite number, providing a viability readout well suited for large-scale drug interaction studies. In contrast, the lower sensitivity of the [^3^H]-hypoxanthine readout precludes similarly sensitive detection when applying a calibration-based approach to relate signal intensity to parasite number after a 120 h recovery period. Comparison of reduced PRR assay with a 7-day recovery period demonstrated that, unlike chemiluminescence, [^3^H]-hypoxanthine incorporation lacks sufficient sensitivity to reliably detect viable parasites, even after 7 days of recovery (Hellingman et al., 2026). Sensitive detection of viable parasites after drug exposure requires at least 14 days of parasite recovery time if the [^3^H]-hypoxanthine incorporation readout is performed (Walz et al., 2023).

Using the MULT-i^2^ assay, we reproduced the time-killing profiles of four reference antimalarials. Interestingly, differences between MULT-i^2^ - and PRR v2-derived profiles were observed for the slower-acting drugs atovaquone, partly also for pyrimethamine, in the characterization of the lag phase and/or the number of viable parasites at later time points, affecting parameters such as the log_10_(PRR). Higher viable parasite counts detected at later time points in the MULT-i^2^ assay may reflect reporter enzyme accumulation during the 120 h recovery window. In contrast, the PRR v2 excludes parasites that die during the 2-week recovery period, since measurements are restricted to a 24 h endpoint window and the readout is based solely on the incorporation of tritium by growing and replicating parasites. Minor signal contributions from sexual parasite stages and mature gametocytes may also elevate the MULT-i^2^ assay signal, as these non-replicating parasite stages can contribute to the chemiluminescence readout in contrast to the PRR v2 assay (Lucantoni et al., 2015; Reader et al., 2015). Although, the use of asynchronous cultures minimizes the impact of stage-specific effects by providing a mixed parasite population representative of the natural distribution of developmental stages, differences in parasite stage progression following drug exposure may still contribute to variation in the extrapolated parasite numbers. Furthermore, compounds that alter parasite metabolism or induce delayed recovery or transient drug-induced dormancy may influence parasite proliferation after treatment, potentially biasing the inferred parasite numbers. The underlying assumption of the linear regression, relating signal intensity to parasite number, is that enzyme accumulation in treated parasites mirrors that in untreated parasites. Deviations from this assumption can lead to over- or underestimation of parasite numbers, potentially explaining the shorter lag phase observed for atovaquone in the MULT-i^2^ assay compared to the PRR v2 assay. Future optimization of rapamycin exposure duration may improve model fit by mitigating artifacts in overgrown cultures. Here, decrease in luminescence following prolonged incubation likely reflects reporter enzyme degradation rather than true parasite loss.

In the dual-drug combination experiments we observed comparable interaction patterns between MULT-i^2^ and combination PRR (cPRR) assays. Importantly, the MULT-i^2^ assay enabled testing of 49 drug combinations compared to only 9 combinations that can be tested with the conventional cPRR assay. Thus, the rich MULT-i^2^ assay dataset supported more robust GPDI parameter estimation than cPRR assay data, evidenced by smaller relative standard errors of estimated parameters and residual errors. The rich dataset obtained by the MUTL-i^2^ assay, spanning a much bigger concentration range, also enabled the identification of additional significant interactions.

For the validation of the well-known synergistic combination of atovaquone and proguanil, both the MULT-i^2^ and cPRR assay revealed the same dominant interaction, with proguanil reducing the EC50 of atovaquone. This is consistent with published notions that proguanil enhances the effect of atovaquone on collapsing the mitochondrial membrane potential (Srivastava Indresh & Vaidya Akhil, 1999). For pyronaridine and piperaquine, the interaction is synergistic at low concentrations but shifts toward antagonism at higher concentrations, as observed with both assays. This concentration-dependent pattern may be partially explained by increased competition for shared targets or reduced target accessibility at elevated drug levels, given that both drugs act within the digestive vacuole and inhibit hemozoin formation (Croft et al., 2012; Dhingra Satish et al., 2017; Herraiz et al., 2019; Warhurst et al., 2007). Additionally, deviations from the Bliss Independence null-interaction model may indicate that full pharmacological independence, an assumption underlying Bliss Independence (Pearson et al., 2023; Roell et al., 2017), is not achieved. This may be due to the partially overlapping mechanisms of action of the two drugs.

One limitation of the MULT-i^2^ assay is its reliance on transgenic *P. falciparum* parasites. To test field isolates at a larger scale than is manageable with the cPRR assay, alternative readouts, such as MitoTracker-based assays (Maiga et al., 2024; Maiga et al., 2025), have been deployed, but their throughput is limited due to the flow cytometry-based readout. Another approach is to compute and predict optimized experimental design layouts (Chen et al., 2018; Kroemer et al., 2022) for the experiments to reduce workload and maximize information gained with the conventional cPRR viability readout.

The dependence on transgenic parasites notwithstanding, the MULT-i^2^ assay using a non-radioactive readout, represents a substantial advance for systematic *in vitro* antimalarial combination testing. Compared to the cPRR assay, it reduces resource requirements by >50-fold and assay duration by more than twofold, enabling the exploration of much larger interaction spaces. The resulting, data-rich datasets support more precise pharmacodynamic modeling and facilitate the identification of previously unrecognized interactions, while improving overall cost-effectiveness. Here we demonstrate dual-drug testing, but the approach is readily extendable to triple combinations, which is important to inform future preclinical and clinical studies, as systematic data on triple antimalarial combinations, including triple artemisinin combination therapies (Hamaluba et al., 2021; van der Pluijm et al., 2019; van der Pluijm et al., 2020), remain limited. In addition, the readout to quantify parasite viability no longer relies on radioactivity or expensive devices, such as flow cytometers. Instead, a standard plate reader capable of detecting luminescence is sufficient.

In conclusion, the non-radioactive MULT-i^2^ assay provides a scalable, sensitive, and cost-effective platform for studying single- and dual-drug combinations, while reducing experimental resource requirements by more than an order of magnitude and shortening assay time. It enables high-resolution mapping of pharmacodynamic interactions and is readily extendable to triple-drug combinations, thereby advancing antimalarial pharmacodynamic modeling and supporting the systematic development of combination therapies.

## Material and Methods

### P. falciparum cultivation

We cultivated parasites using standard methods (Haynes et al., 1976; Hellingman et al., 2024; Snyder et al., 2007; Trager & Jensen, 1976). Culture medium (CM) consisted of RPMI 1640 (10.44 g/L, Thermo Fisher Scientific, Massachusetts, USA), HEPES (5.94 g/L, Sigma-Aldrich, Buchs SG, Switzerland), NaHCO_3_ (2.1 g/L, Sigma-Aldrich), neomycin (100 mg/ L, Sigma-Aldrich), Albumax II (5 g/L, Thermo Fisher), and hypoxanthine (50 mg/L, Sigma-Aldrich). All components were dissolved in water (Milli-Q purified), and the CM was sterile-filtered using a bottle top filter (0.22 μm pore size, Corning, New York, USA). Parasite cultures were kept at a hematocrit (hc) of 5% and incubated at 37 °C in a malaria gas mixture consisting of 93% N_2_, 4% CO_2_, and 3% O_2_. Erythrocyte concentrates were obtained from the local blood bank.

### Generation of transgenic P. falciparum line with inducible lacZ expression

Transfection of *P. falciparum* NF54^DiCre^ parasites (Tibúrcio et al., 2019) was performed using standard protocols developed by Voss et al. (Voss et al., 2006). Genome editing of the *P230p* locus (PF3D7_0208900) was achieved by co-transfecting the pHF-gC-*P230p* plasmid, carrying Cas9 and a *P230p*-targeting guideRNA (Ashdown et al., 2020; Ghorbal et al., 2014), with the *P230p*-loxPIntron-GFP-*lacZ* rescue plasmid (codon optimized *E. coli lacZ*), derived from the previously described *P230p-lacZ* plasmid (Hellingman et al., 2024). For positive selection, parasites were cultivated with 5 nM of the antifolate drug WR99210, selecting for the human dihydrofolate reductase (hDHFR) marker present on pHF-gC-*P230p*. Negative selection was then applied using 10 µM of the prodrug 5-fluorocytosine (5-FC), selecting against the same plasmid via the yeast cytosine deaminase and uridyl-phosphoribosyltransferase (yFCU) marker (Braks et al., 2006; Manzoni et al., 2014). To obtain clones of the *^i-lacZ^*, limiting dilution was carried out in a 96-well plate, and clone T30-E6 was selected for subsequent experiments. Site-specific recombination was induced using a non-growth-altering concentration of 100 nM rapamycin (Sigma-Aldrich) (Atack et al., 2020; Knuepfer et al., 2017; Veletzky et al., 2014). Diagnostic integration PCR (KAPA HiFi HotStar PCR Kit, Kapa Biosystems, Roche, Massachusetts, USA) was used to verify the correct editing of the *P230p* locus (details regarding the diagnostic PCR thermocycler program can be found in Supplementary Table 1). The primer sequences were as follows: #1 5’-AGACACACCATCAACATTATCG, #2 5’-TTGTAAAACTTGGACAAGCTAAACC, #3 5’-GAAAATCCTAAATTATGGAGTG and #4 5’-CCAGTCTCTTCTCTGCAGG. For marker control (hDHFR and1.2 yFCU; Figure 1 – figure supplement 1): #5 5’-GACCTCTGTCGAGGGC and #6 5’-GTCCCAGAACATGGGC. PCR positive control (1252 bp) primers were targeting the *mdr1* locus (PF3D7_0523000) with forward primer 5’-ATGGGTAAAGAGCAGAAAGAG and revers primer 5’-CACAACCTGATTCTCCCACAAAT (Bravo et al., 2025). As DNA ladder a 1 kb ladder with size range from 0.5 to 10 kb (New England Biolabs, Massachusetts, USA) was used.

### Recombination efficacy of the P. falciparum ^i-lacZ^ line

We adjusted asynchronous parasites to ∼0.3% parasitemia and cultured at 1.25% hc under three conditions: with 100 nM rapamycin, dimethyl sulfoxide (DMSO, Sigma-Aldrich), or no treatment. After 48 h, samples were stained with 5 μg/mL Hoechst 33342 (Thermo Fisher Scientific) diluted in phosphate-buffered saline (PBS) for 20 min, washed twice with PBS, and imaged by fluorescence microscopy (DM5000B, 100-fold objective, Leica, Wetzlar, Germany). GFP and Hoechst signals were recorded using Leica Application Suite X (version 3.7.5.24914), and images were analyzed with ImageJ (version 2.16.0/1.54p/Java1.8.0_322 (64-bit)). To assess *lacZ* expression, additional parasite culture samples treated with 100 nM rapamycin or DMSO for 48 h were taken. Uninfected RBCs (uRBCs) at the same hematocrit, treated with 100 nM rapamycin, served as negative control. Samples were distributed into white flat-bottom 96-well plates (Greiner Bio-One, Kremsmünster, Austria) and incubated with the β-gal^SENSOR^ probe (AquaSpark β-D-galactoside, cat. #A-8169_P00, Biosynth AG, Staad, Switzerland) dissolved in ddH₂O containing MgCl₂, to final concentrations of 10 µM β-gal^SENSOR^ probe and 200 µM MgCl₂. Plates were incubated at 37 °C for 30–60 min, and luminescence (counts/s) was measured using a Spark multimode reader (TECAN, Männedorf, Switzerland) at 37 °C with a 5 s exposure time.

We analyzed the same parasite cultures also by High Content Imaging (HCI). Hoechst-stained samples were diluted to ∼0.02% hc, and 200 µL was transferred into black µClear 96-well plates (Greiner Bio-One). Cells were allowed to settle for ∼15 min before imaging. Fluorescence microscopy images (9×9 raster; 81 images per well) were acquired for GFP and Hoechst using the ImageXpress Micro XLS high content imager (Molecular Devices, California, USA). Hoechst- and GFP-positive cells were quantified with automated image analysis (MetaXpress, 64-bit, version 6.7.1.157). Detailed image analysis parameters are provided in Supplementary Table 2. Recombination efficacy after 48 h of rapamycin treatment was calculated as follows:

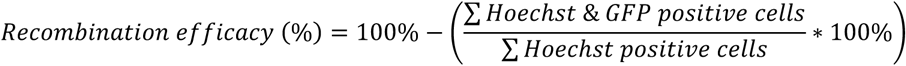

Values represent the average of three biological replicates (experiment(s) conducted with independent sample(s)).

### Parasitized erythrocyte infection rate

The infection rate of erythrocytes by the transgenic *P. falciparum* strain *^i-lacZ^*, with or without induction by 100 nM rapamycin, was compared to that of the wild type NF54 strain using mixed-stage parasite cultures. Parasite multiplication rates were calculated from parasitemia determined at 0 h (initial parasitemia ∼0.3%) and after 48 h, based on counts from Giemsa-stained blood smears. Cultures were maintained at a hematocrit of 1.25% under standard cultivations conditions.

### Evaluation of limit of quantification

To evaluate the parasite limit of detection under different incubation time periods following treatment with 100 nM rapamycin, the number of asynchronous *P. falciparum ^i-lacZ^* was set to 750’000 per mL. This was calculated based on parasitemia determined with Giemsa-stained blood smears and the number of RBCs per mL quantified with an Improved Neubauer hemocytometer. Parasites were dispensed in technical duplicates into white, flat-bottom 96-well plates and serially diluted two-fold resulting in a theoretical parasite range of 75’000 – 0.14 parasites per well. Rapamycin or DMSO (vehicle control) was diluted in culture medium and added to the wells to a final concentration of 100 nM. As negative controls, uninfected RBCs also treated with 100 nM rapamycin or DMSO were included. The final hematocrit was 1.25%. The 96-well plates were then incubated for 24, 48, 72, 96, 120, 144, or 168 hours in humidified gas chambers under cultivation conditions before they were frozen at −20°C. For analysis, plates were thawed, and the β-gal^SENSOR^ probe, dissolved in ddH_2_O containing MgCl_2_, was added to each well to reach final concentrations of 10 µM β-gal^SENSOR^ probe and 200 µM MgCl_2_. Plates were then incubated at 37°C for 30-60 min, and luminescence (counts/s) was measured with the Spark multimode reader (TECAN) at 37°C with an exposure time of 5 s. Experimental data were analyzed using GraphPad Prism (version 10.0.1). The limit of quantification (LOQ) of the chemiluminescence signal from transgenic parasites was determined using an unpaired t-test between the signal from each parasite dilution and the negative control (uRBC treated with rapamycin), along with linear regression of log_10_(parasites/well) versus log_10_(measured counts/s). The smallest number of parasites per well for which the t-test was considered significant (p < 0.05) and a linear relationship between log_10_(parasites/well) vs. log_10_(measured counts/s) was observed was considered as the parasite limit of quantification for that incubation period under rapamycin treatment.

### IC_50_ assay - [^3^H]-hypoxanthine incorporation

We dissolved compounds in DMSO or sterile filtered purified water (for chloroquine) to obtain a final drug stock concentration of 10 mM. Drug stock solutions were then validated by the [^3^H]-hypoxanthine incorporation inhibitory concentration (IC_50_) assay according to the protocol of Snyder et al. (Snyder et al., 2007). In brief, two-fold serial dilutions of the compounds were performed in 96-well plates in hypoxanthine deficient medium. Infected red blood cells were added resulting in a final hematocrit of 1.25% and parasitemia of 0.3%. The 96-well plates were incubated for 48 hours in humidified gas chambers under standard *P. falciparum* cultivation conditions. After this period, [^3^H]-hypoxanthine (American Radiolabeled Chemicals, Missouri, USA) was added, and the plates were incubated for an additional 24 h. To quantify incorporated [^3^H], plate contents were harvested onto glass fiber filter mats, and signal was measured using a Betaplate^TM^ liquid scintillation counter (PerkinElmer, Massachusetts, USA). IC_50_ values were calculated by linear interpolation (Huber & Koella, 1993; Snyder et al., 2007). For drug stock validation, calculated IC_50_ values were then validated according to published IC50 reference values of the respective compounds for NF54 (artemisinin 5.8 nM (SD 2.3); chloroquine 11.2 nM (SD 2.7); pyrimethamine 17 nM (SD 8.0); atovaquone 0.5 nM (SD 0.1); proguanil 1115.3 nM (SD 712.0); pyronaridine 4.9 nM (SD 0.8); piperaquine 8.3 nM (SD 1.0)) (Delves et al., 2012).

### Multidimensional luminescence test for integration of interactions (MULT-i^2^) assay

The developed protocol to test the viability of the transgenic *P. falciparum ^i-lacZ^* after single- or combination-drug treatment was performed as follows: Based on parasitemia and RBCs per mL (determined using an Improved Neubauer hemocytometer), an infected RBC (iRBC) suspension containing 10^6^ asynchronous parasites/mL at a hematocrit of 2.5% was prepared (corresponding to a parasitemia of ∼0.4%). Drug working solutions were diluted in culture medium using a 10 mM drug stock solution in DMSO.

For single-drug testing, 100 uL of each drug working solution of four reference antimalarials - artemisinin, chloroquine, pyrimethamine and atovaquone - was distributed in technical quadruplicates across five white, flat-bottom 96-well plates (one plate per tested time point). Subsequently, 100 uL of the prepared iRBC suspension was added to each well, resulting in a theoretical parasite number of 10^5^ per well and a final hematocrit of 1.25% (corresponding to a parasitemia of ∼0.4%). The final tested drug concentrations corresponded to 10×IC_50_ values (with atovaquone as an exception): artemisinin (58 nM), chloroquine (112 nM), pyrimethamine (170 nM) and atovaquone (100 nM). As controls, uninfected RBCs (uRBCs) and untreated iRBCs were included on each plate.

For dual-drug combination testing, drug working solutions were prepared in 96-deep-well (2 mL) plates (Greiner Bio-One). Two-fold (for pyronaridine-piperaquine) or three-fold (for atovaquone-proguanil) checkerboard dilutions were performed to obtain 7×7 different concentration combinations and seven single-drug concentrations per tested compound (Bellio et al., 2021). A 100 uL aliquot of each prepared working drug dilution was transferred to five white, flat-bottom 96-well plates in one technical replicate (measurement(s) of same sample in biological replicate). Then, 100 uL of the prepared iRBC suspension was added to each well, resulting in a theoretical parasite number of 10^5^ per well and a final hematocrit of 1.25%. Final tested drug concentrations were as follows: for pyronaridine 50, 25, 12.5, 6.25, 3.125, 1.563, 0.781 nM; for piperaquine 200, 100, 50, 25, 12.5, 6.25, 3.125 nM; for atovaquone 300, 100, 33.3, 11.1, 3.7, 1.23, 0.41 nM and for proguanil 50’000, 16’666.7, 5’555.6, 1’851.9, 617.3, 205.8, 68.6 nM. In addition, all the 49 possible dual-concentration combinations between pyronaridine + piperaquine and atovaquone + proguanil were tested. As controls, uRBCs, and untreated iRBCs were included on each 96-well plate.

For each drug testing experiment, we prepared an additional 0 h reference calibration plate (white, flat-bottom plate) using a two-fold serial dilution of the parasite suspension, starting from 10^5^ to 0.19 parasites per well, with a final hematocrit of 1.25%. For parasite growth controls, untreated parasite suspensions were distributed into transparent 6-well plates (Falcon) with a final hematocrit of 1.25%.

All prepared plates (drug-treated plates, 0 h reference calibration plates, and growth control plates) were incubated in a humidified gas chamber under standard cultivation conditions. Drug and CM of the single-drug testing experiments were replaced every 24 h. After the desired incubation period under drug pressure (0 h for the reference calibration plate; 24 h, 48 h, 72 h, 96 h and 120 h for drug-treated plates), the iRBC pellet was washed at least three times for single-drug testing and five times for combination testing to remove the drug (the reference calibration plate was washed identically for procedural consistency). The washing procedure consisted of repeated rounds of supernatant removal, addition of fresh culture medium, and centrifugation of the 96-well plates at 600g for 2 min. After the final centrifugation step of the 24 h plate, the supernatant was collected for washout control analysis (for dual-combination assays, samples were collected from the wells containing the two highest single-drug concentrations and the highest dual-drug combination.) Following drug washout, rapamycin (final concentration 100 nM) was added to all drug-treated wells, uRBCs (negative control), and one subset of untreated iRBCs (positive control). To the other subset of untreated iRBCs, DMSO was added (negative iRBCs control). After rapamycin or DMSO addition, plates were incubated for 120 h and then frozen at −20 °C until measurement. For the growth control cultures (prepared in the 6-well plate), parasitemia was determined from three independent Giemsa-stained blood smears, each based on counts of 10’000 RBCs at 0 h and after 48 h of incubation.

For analysis, the 96-well plates were thawed, and the β-gal^SENSOR^ probe (dissolved in ddH_2_O containing MgCl_2_) was added to each well to achieve final concentrations of 10 µM β-gal^SENSOR^ probe and 200 µM MgCl_2_. The plates were then incubated at 37 °C for 30-60 min, and luminescence (counts/s) was measured using a Spark multimode reader (TECAN) at 37 °C with an exposure time of 5 s.

### MULT-i^2^ assay data processing for modeling

We used the 0 h reference calibration plate to establish the linear relationship between the initial parasite number per well and the corresponding chemiluminescent signal intensity. A linear regression was therefore performed between log_10_(initial parasites per well) and log_10_(measured counts/s) for the initial parasite number datapoints that showed a significant difference (p < 0.05) from the negative control (uRBC treated with Rapa) in an unpaired t-test, and that showed a linear regression equation with a R square value ≥0.99:

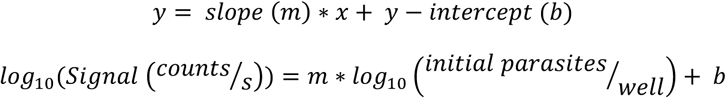

The calculated slope (m) and y-intercept (b) from the linear regression were used to determine the initial parasite number per well based on the measured luminescent signal intensity. The resulting values were then normalized (to correspond to 10^5^ parasites at 0 h), and the mean plus three standard deviations (mean + 3*SD) of the uRBC negative control signal was subtracted to correct for background luminescence. If previous time points of the same tested concentration were negative for parasite viability, the following time points were also set negative for parasite viability. The parasite growth rate was calculated from parasitemia values counted at 0 h and 48 h. Data processing and visualization were performed using GraphPad Prism (version 10.0.1), Excel (version 2507 Build 16.0.19029.20136) 64-bit), R (version 4.5.1 (2025-06-13 ucrt)), and RStudio (version 2025.05.1+513)

### Drug washout control analysis

To assess the efficiency of drug washout, we collected supernatant after the final centrifugation step of the washout process and diluted it in CM following the same procedure used for the experimental samples (adjusted to a 2-fold higher concentration to account subsequent addition of parasite suspension) as proposed by Walz et al. 2023. In brief, NF54^i-lacZ^ parasites were then exposed, in technical duplicates, to a serial dilution of this collected an diluted supernatant at 1.25% hematocrit and 0.3% parasitemia and incubated at 37 °C under growth conditions. Parasite growth was assessed by incorporation of tritium-labelled hypoxanthine added 48 h after initiation, followed by plate freezing at −20 °C after an additional 24 h and harvesting of lysed cells as described in the section “IC_50_ assay - [^3^H]-hypoxanthine incorporation”. Drug washout was considered successful when parasite growth in the washout control samples was comparable to that of the positive control (untreated parasites), defined as less than 20% deviation between the two.

### Combination PRR (cPRR) assay

The cPRR assay, performed to compare results with the MULT-i^2^ assay, was adapted from Sanz et al. (Sanz et al., 2012). In brief, we adjusted asynchronous *P. falciparum* parasites (NF54) to a hematocrit of 2% and parasitemia of 0.5% (with ≥60% ring stages) and distributed into 6-well plates. Cultures were incubated in humidified gas chambers under standard cultivation conditions, either in CM alone (growth control) or under drug pressure. Tested drug concentrations were as follows: Atovaquone 0.5, 3, 10 and 100 nM (3 nM excluded from combo testing) and proguanil 70, 1500, 16000 and 50000 nM (16000 nM excluded from combo testing). All nine possible dual combinations between atovaquone and proguanil were included. Pyronaridine 3, 5, 6 and 25 nM (5 nM excluded from combo testing) and piperaquine 8, 10, 12 and 100 nM (10 nM excluded from combo testing). All nine possible dual combinations between pyronaridine and piperaquine were included. As an assay control, the reference compound pyrimethamine was teste at 10×IC_50_ (229 nM) for 24, 48, 72 and 96 hours.

For atovaquone + proguanil, aliquots (1 mL) were collected at time points 48, 72 and 120 h, and for pyronaridine + piperaquine, at 24, 48 and 120 h. Drugs were removed by three washing cycles consisting of centrifugation (600 g, 1 min), supernatant removal and addition of 1 mL fresh CM. For pyrimethamine-treated cultures, the drug and CM were replaced every 24 h, and samples were collected accordingly. At each time point, Giemsa-stained blood smears were prepared and parasitemia was determined microscopically. Washed aliquots were distributed into transparent 96-well plates (Sarstedt) in four technical replicates (and eight replicates for time point 0 h). Samples were serially diluted 15 times with a 2.8 dilution, yielding a final hematocrit of 0.8%. Plates were incubated for 21 days, with weekly medium replacement and addition of fresh erythrocytes at 1% hematocrit. After 20 days of incubation, CM was replaced with 0.5 µCi [^3^H]-hypoxanthine in hypoxanthine-deficient medium, and plates were incubated for an additional 48 h, then frozen at −20 ◦C.

For signal measurement, plates were thawed, and the contents of each well were harvested onto glass fiber filters using a Betaplate™ cell harvester (Perkin Elmer). Filter discoloration was noted. Dried filters were placed in plastic foils containing 3.5 mL scintillation fluid, and radioactivity (counts per minute) was measured using a Betaplate^TM^ liquid scintillation counter (Perkin Elmer). Data processing was performed with Excel (version 2507 Build 16.0.19029.20136) 64-bit).

### Interaction modeling

*In vitro* assay data obtained from the cPRR assay (one biological replicate) or the MULT-i^2^ assay (three biological replicates) were used to build the model (compartmental representation in Supplementary Figure 1). We performed data analysis using nonlinear mixed-effects modeling implemented in the Nonlinear Mixed Effects Modelling (NONMEM) ® software (version 7.5; ICON Development Solutions, Ellicott City, MD, USA). We assumed drug concentrations to remain constant throughout the incubation period. Initially, an exponential growth model was applied to estimate parasite growth parameters, as follows:

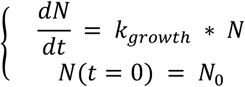

N(t) represents the model-predicted parasitemia at timepoint *t*, N_0_ is the initial parasitemia determined at 0 h, and k_growth_ is the first-order parasite growth rate, estimated from N_0_ and N counted at 48 h in untreated control cultures. A combined proportional-additive residual error model was used to relate observed assay results (Y) to the model predicted parasitemia (N):

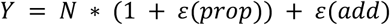

For the atovaquone-proguanil combination the additive error (ɛ(add)) was fixed to 0. For the pyronaridine-piperaquine combination, both, the proportional error (ɛ(prop)) and ɛ(add) were estimated. Values below the lower limit of quantification (LLOQ) were handled with the M3 method (Beal, 2001; Bergstrand & Karlsson, 2009). In the MULT-i^2^ assay, an upper limit of quantification (ULOQ) was introduced because assay signals plateau when parasites overgrow, such as in growth controls or at low drug concentrations. This limit was therefore estimated, and predicted parasitemia values exceeding the ULOQ were capped at N = ULOQ. Next, single-drug effects were estimated from the single-drug assay data using a sigmoidal maximum effect model (Holford & Sheiner, 1981). In this model, drug effect (E) is described as a function of drug concentration (C), maximal drug effect (Emax), half-maximal effective concentration (EC50), and the Hill coefficient (H), which determines the steepness of the concentration-response curve. If a tested drug exhibited a lag phase, a first-order time-delay rate constant (1 - *e*^−*k*^*^lag^*^∗t^) was incorporated into the corresponding killing rate term to account for the delayed onset of drug action:

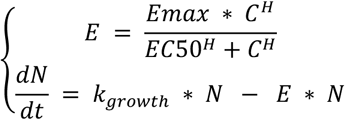

As a third step, the combined drug effects (Ecomb) of compounds A and B were implemented using the Bliss Independence model (Bliss, 1939). The single-drug effects were normalized to 1 to calculate the probabilistic Bliss Independence term, and the resulting values were then rescaled to the original effect scale. For subsequent evaluation, drug null-interaction was assessed under the assumption of no pharmacodynamic interaction, by fixing all GPDI model parameters to zero to obtain pure Bliss Independence.

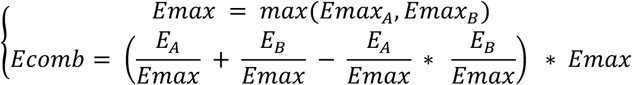

To quantify potential pharmacodynamic interactions, the General Pharmacodynamic Interaction (GPDI) model was applied (Wicha et al., 2017). In brief, shifts in a pharmacodynamic parameter (θ) of the victim drug, induced by the perpetrator drug at concentration C, were incorporated into the drug effect model through the GPDI term. Fractional changes in the pharmacodynamic parameter (θ – either EC50 or Emax) were characterized by the interaction intensity (INT), interaction potency (EC50_INT_) and steepness of the interaction (H_INT_). These three parameters are directional, and when the GPDI term is applied to both tested drug candidates, the interaction becomes bidirectional, indicating that each compound can act as both perpetrator and victim (notation example in the equation below: _AB_ denotes drug A as victim and drug B as perpetrator). When the GPDI term is applied on EC50, the interaction represents a competitive mechanism between drugs A and B (example equation for E_A_). Conversely, when the GPDI term is applied to Emax, it describes an allosteric interaction between drugs A and B (example equation for E_B_):

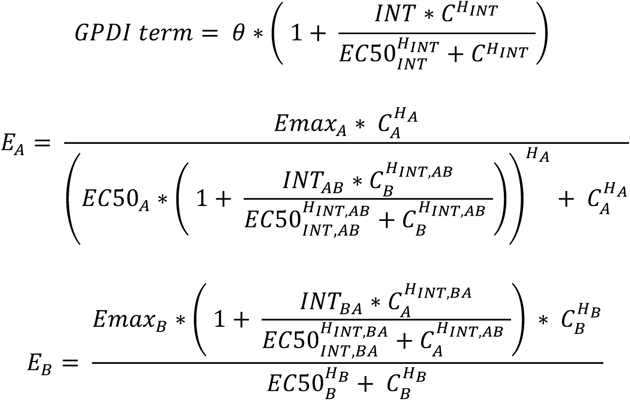

The polarity of the INT parameter, and whether it is applied to EC50 or Emax, determines the type and direction of interaction between the two drugs. An INT=0 indicates no interaction, as the GPDI term equals 1. When the interaction is implemented on EC50, INT values between −1 and 0 (−1<INT<0) describe a synergistic interaction (reflecting a decrease in EC50), whereas INT>0 indicates an antagonistic interaction (increase in EC50). The same polarity of both INT parameters suggests bidirectional synergy or antagonism, while opposite polarities indicate an asymmetric interaction between the two drugs. When the GPDI term is applied to Emax the interpretation of INT polarity is reversed: INT>0 indicates synergism (increase in Emax) and −1<INT<0 denotes antagonism (decrease in Emax). For simplicity, all H_INT_ values were fixed to 1.

The model was constructed stepwise using statistical criteria (likelihood ratio test, α = 0.05, df=1), beginning with a reduced model containing a single INT parameter and its corresponding EC50_INT_, and progressively evaluating all possible INT parameters with both drugs acting as perpetrator and/or victim. Model fit was assessed through graphical visualization and the Akaike information criterion (AIC), whereas a lower AIC value relative to the null-interaction model indicated an improved fit, which was then selected for further analysis. Example .mod files for modeling can be found in the supporting materials.

## Supporting information

genome editing strategy

IC50 analysis

compartmental representation scheme

## Additional information

### Author contributions

Angela Hellingman, Idea and conceptualization, Conducting and validating *in vitro* experiments, Conducting and validating *in silico* modeling, Writing - original draft preparation, Writing – review and editing; Christin Gumpp, Conducting and validating *in vitro* laboratory experiments, Writing – review and editing; Jörg J. Möhrle, Idea and conceptualization, Supervision, Writing – review and editing, Project administration and funding acquisition; Belen Tornesi, Writing – review and editing, Project administration and funding acquisition; Didier Leroy, Writing – review and editing, Project administration and funding acquisition; Sergio Wittlin, Idea and conceptualization, Supervision, Writing – review and editing, Project administration and funding acquisition; Pascal Mäser, Supervision, Writing – review and editing, Project administration and funding acquisition; Nicolas M. B. Brancucci, Idea and conceptualization, Supervision, Writing – review and editing; Sebastian G. Wicha, Idea and conceptualization, Conducting and validating *in silico* modeling, Supervision, Writing – review and editing; Matthias Rottmann, Idea and conceptualization, Supervision, Writing – review and editing, Project administration and funding acquisition.

## Acknowledgments

The authors would like to thank Biosynth AG for offering a discount on the AquaSpark β-D-galactoside probe.

## Abbreviations

ACTs: artemisinin-based combination therapies
PRR: parasite reduction ratio
PCT: parasite clearance time
EC50: half-maximal effective concentration
Emax: maximal drug effect
HRP-2 ELISA: histidine-rich protein 2 sandwich enzyme-linked immunosorbent assay
MULT-i^2^: multidimensional luminescence test for integration of interactions
GPDI: general pharmacodynamic interaction model
*gfp* or GFP: green fluorescent protein
*hsp70*: *heat shock protein 70*
*cam*: *calmodulin*
loxPint: loxP-intron
DMSO: dimethyl sulfoxide
uRBC: uninfected red blood cells
iRBC: infected red blood cells
rapa: rapamycin
GFP: green fluorescent protein
LOQ: limit of quantification
PRR v2: parasite reduction ratio version 2 assay published by Walz et al.
cPRR: combination PRR assay
AIC: Akaike Information Criterion
NONMEM: Nonlinear Mixed Effects Modelling
DV: dependent variable
IPRED: individual predicted values
CM: culture medium
E: effect
C: concentration
Ecomb: combination effect
H: hill coefficient
INT: interaction parameter
EC50_INT_: half-maximal effective concentration of interaction
H_INT_: hill coefficient
IC_50_: half-maximal inhibitory concentration
LLOQ: lower limit of quantification
ULOQ: upper limit of quantification.

## Additional files

**Supplementary file 1:** Details about the used diagnostic PCR program, and the automated HCI analysis for calculation of recombination efficacy; detailed table of model estimates; compartmental representation of the developed pharmacometric model; example .mod files for Bliss Independence – null-interaction or with interactions.

**Supplementary figure 1:** Compartmental representation of the developed pharmacometric model. Assay signal was translated to parasitemia. Drug effects were modelled using a sigmoidal maximum effect model and Bliss Independence was used as null-interaction model with Emax scaled. For atovaquone, drug effect was modeled with a lag time which is implemented on Emax with the term (1-e^−klag×^ ^t^). A, compound A; B, compound B; N0: initial parasitemia, kg: growth rate, Emax: maximal drug effect - maximum killing rate, EC50: concentration stimulating 50% of Emax, Hill: Steepness of the concentration-effect relationship, klag: first-order delay rate for atovaquone until full killing rate is achieved.

**Figure 1B – source data 1:** Original agarose gel image of diagnostic PCRs confirming the integration of the loxPInt-*gfp*-*lacZ* expression cassette in *P. falciparum* NF54*^i-lacZ^*, successful recombination after DiCre-induced recombination, and loss of selectable markers in the NF54*^i-lacZ^* (marker control).

**Figure 1D – source data 2:** Chemiluminescence signal after rapamycin-induced recombination measured 48 hours after rapamycin addition.

**Figure 1 – figure supplement 1:** Overview of the two-plasmid CRISPR/Cas9-based gene editing strategy. Schematic of the *P230p* locus in NF54^DiCre^ parasites and the plasmids used for CRISPR/Cas9-mediated gene editing (*P230p*-loxPInt-gfp-*lacZ* and pHF-gC-*P230p*) to generate the NF54*^i-lacZ^* parasites.

**Figure 1 – figure supplement 2:** Susceptibility to the antifolate drug WR99210 and parasitized erythrocyte infection rate of the novel NF54^i-lacZ^ compared to NF54^WT^. (A) IC_50_ values of WR99210; error bars represent the standard error of the mean (SEM) of three biological replicates conducted in technical duplicates and (B) parasitized erythrocyte infection rates (within a period of 48 hours) are comparable between NF54^i-lacZ^ parasites, either rapamycin-treated or untreated, and the NF54^WT^ strain; error bars represent the SEM of three biological replicates. Rapa: rapamycin.

**Figure 1 – figure supplement 2 – source data 1:** Measured and normalized parasite growth (% of NF54^wt^ or NF54*^i-lacZ^*) versus WR99210 concentration (nM) used for IC50 determination and inhibitory dose–response curve fitting.

**Figure 1 – figure supplement 2 – source data 2:** Microscopically counted parasitemia and proportion of ring stages (%) used to calculate the parasitized erythrocyte infection rate within 48 hours (one asexual intraerythrocytic developmental cycle).

**Figure 2 – source data 1:** Measured chemiluminescence signal for the initial parasite inoculum or the uninfected red blood cell control after the corresponding incubation time under 100 nM rapamycin or DMSO.

**Figure 4 – source data 1:** Dataset of the MULT-i^2^ assay containing the log(viable parasites) per tested time point of the four reference compounds used for plotting time killing profiles and calculating lag time, log_10_(PRR), PCT_99.9%_ and Emax with the R pipeline published by Walz et al., 2023.

**Table 1 – source data 1:** same as indicated in Figure 4 – source data 1

**Figure 5 – source data 1:** Original cPRR dataset used for modeling in the Nonlinear Mixed Effects Modelling (NONMEM) software containing measured values (also called dependent variables (DV), parasitemia %) at tested time points (TIME, hours), used drug concentrations for atovaquone (CA, nM) and proguanil (CB, nM). The original result datasets contain the final plotted model estimates (individual predicted values (IPRED), parasitemia %) for BI null-interaction (orange) or BI Interactions (blue).

**Figure 5 – source data 2:** Original MULT-i^2^ dataset used for modeling in NONMEM containing measured values (DV, parasitemia %) at tested time points (TIME, hours), used drug concentrations for atovaquone (CA, nM) and proguanil (CB, nM). The original result datasets contain the final plotted model estimates (IPRED, parasitemia %) for BI null-interaction (orange) or BI Interactions (blue).

**Figure 6 – source data 1:** Original cPRR dataset used for modeling in NONMEM containing measured values (DV, parasitemia %) at tested time points (TIME, hours), used drug concentrations for pyronaridine (CA, nM) and piperaquine (CB, nM). The original result datasets contain the final plotted model estimates (IPRED, parasitemia %) for BI null-interaction (orange) or BI Interactions (blue).

**Figure 6 – source data 2:** Original MULT-i^2^ dataset used for modeling in NONMEM containing measured values (DV, parasitemia %) at tested time points (TIME, hours), used drug concentrations for pyronaridine (CA, nM) and piperaquine (CB, nM). The original result datasets contain the final plotted model estimates (IPRED, parasitemia %) for BI null-interaction (orange) or BI Interactions (blue).

## Data availability

All data generated or analyzed during this study are provided with the manuscript and its supplementary files, including individual and averaged source datasets used to generate the figures and supplementary figures. Data from Walz et al. (2023), used for comparison in Figure 4, are available from the original publication. Further information is available from the corresponding authors upon request.

## References

### Figures

Figure 3 was partly created with biorender.com, Rottmann, M.

*Tools for language improvement (rephrasing and translating)*

- DeepL Write, Deepl SE: https://www.deepl.com/write; rephrasing of text passages
- ChatGPT, version 5, OpenAI, https://chatgpt.com/; rephrasing of text passages

