## Supplementary figures and images for "Multidimensional *in vitro* assay for antimalarial combination testing and pharmacodynamic modeling – the MULT-i^2^ assay"

### compartmental representation scheme

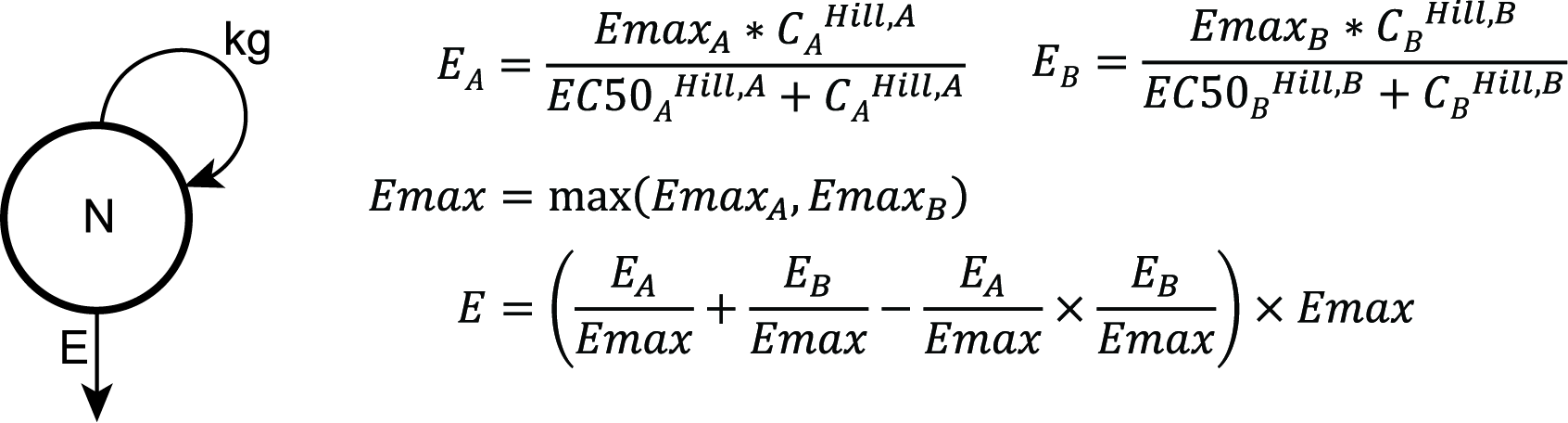

### genome editing strategy

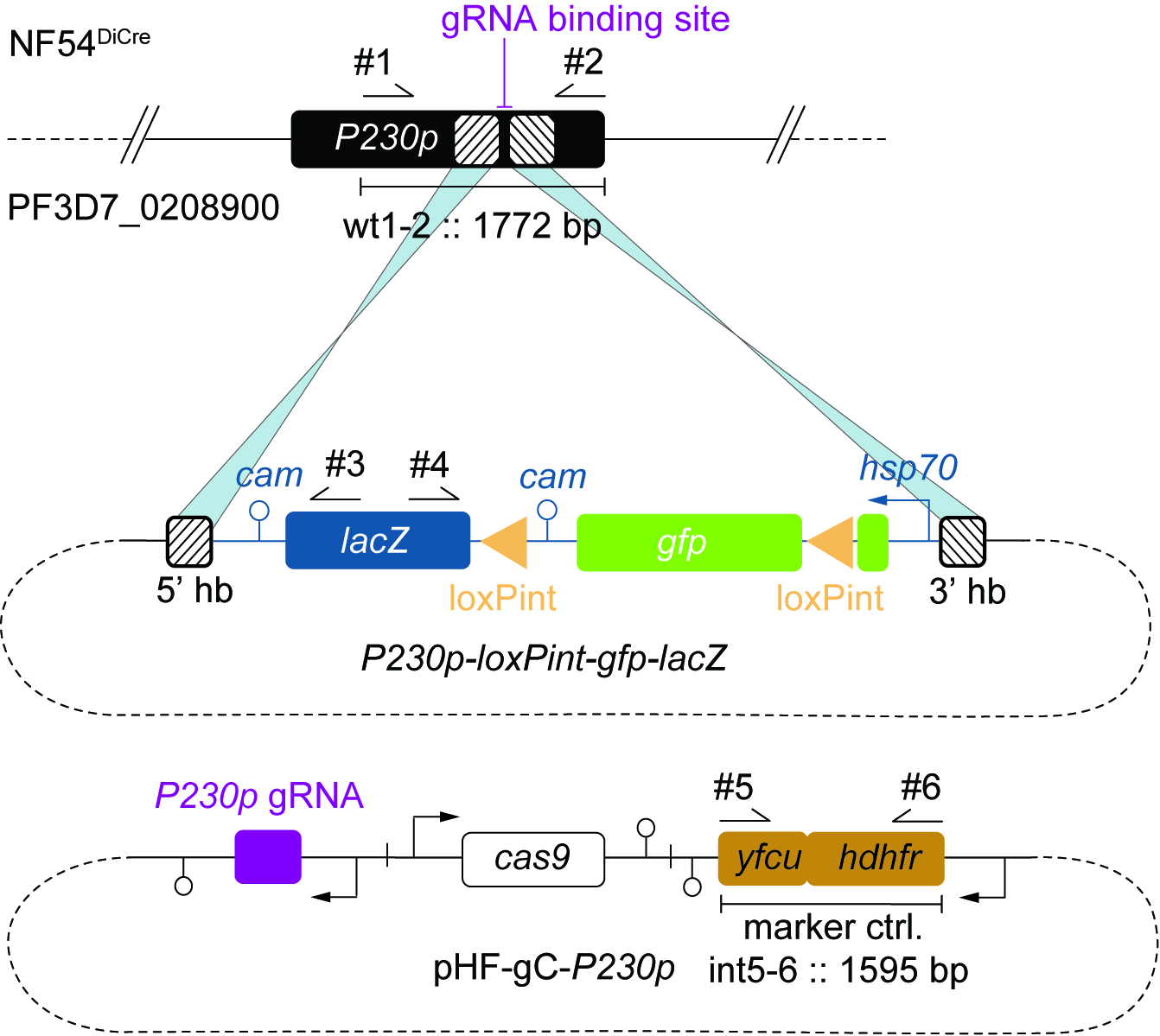

### IC50 analysis

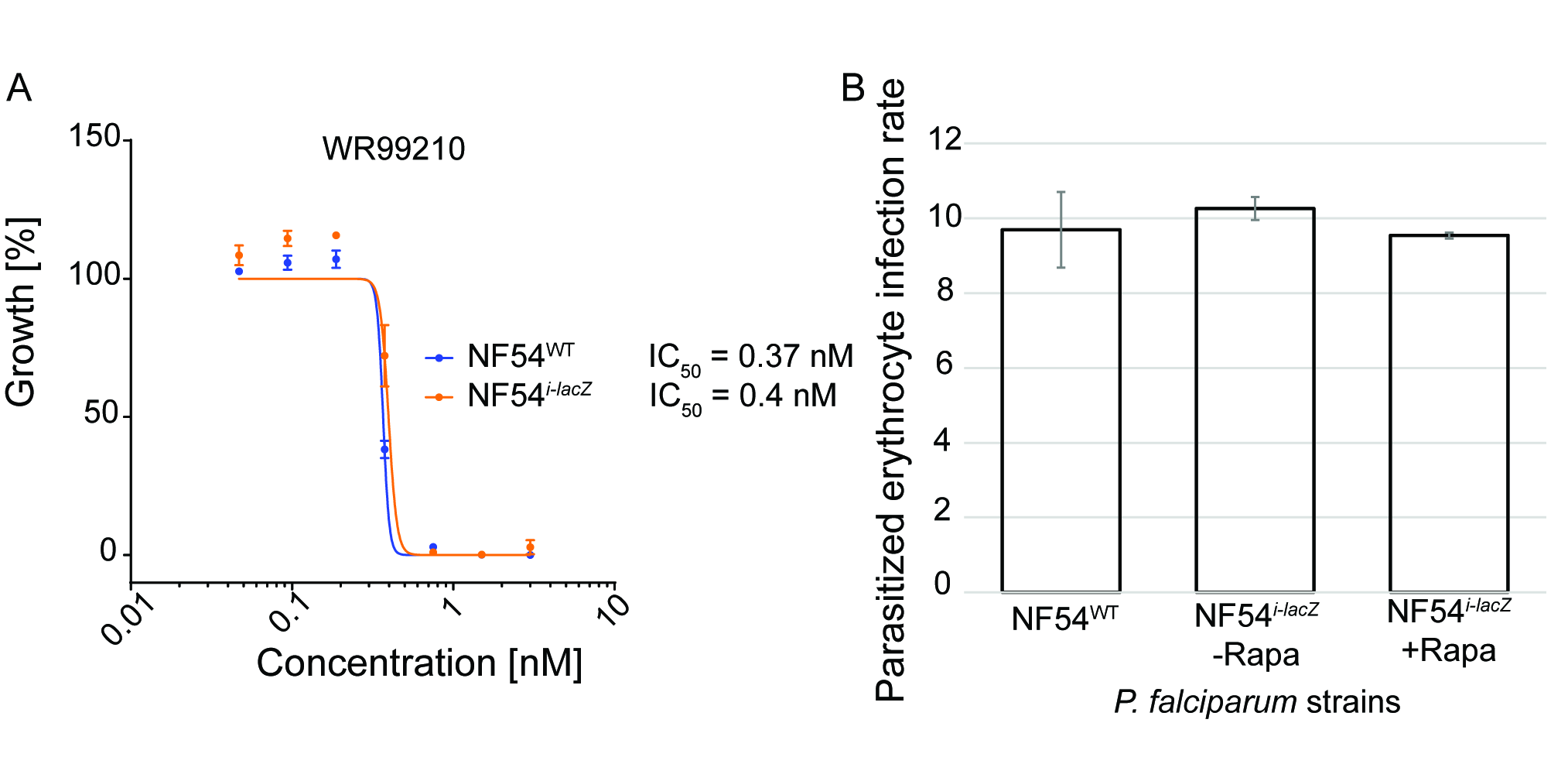
